# NHR-49 Acts in Distinct Tissues to Promote Longevity versus Innate Immunity

**DOI:** 10.1101/2020.09.11.290452

**Authors:** Nikki Naim, Francis RG Amrit, Ramesh Ratnappan, Nicholas DelBuono, Julia A Loose, Arjumand Ghazi

## Abstract

Aging and immunity are inextricably linked and many genes that extend lifespan also enhance immunoresistance. However, it remains unclear if longevity-enhancing factors modulate immunity and longevity by distinct or shared mechanisms. Here, we demonstrate that the *Caenorhabditis elegans* pro-longevity factor, NHR-49, also promotes resistance against *Pseudomonas aeruginosa*, but modulates immunity and longevity by spatially and mechanistically distinct mechanisms. Fenofibrate, an agonist of NHR-49’s mammalian functional homolog, PPARα, enhanced worm immunoresistance in an NHR-49-dependent manner. NHR-49 expression is increased by germline ablation, an intervention that extends lifespan, but lowered by pathogen exposure. NHR-49 acted in multiple somatic tissues to promote longevity, whereas, it’s pro-immunity function was mediated by neuronal expression. The canonical NHR-49 target genes, *acs-2* and *fmo-2*, were upregulated by germline loss, but infection triggered *fmo-2* downregulation and *acs-2* upregulation. Interestingly, neither gene conferred resistance against Gram-negative *Pseudomonas*, unlike their reported roles in immunity against Gram-positive pathogens. Thus, NHR-49 is differentially regulated by interventions that bring about long-term changes (lifespan extension) vs. short-term stress (pathogen exposure) and in response it orchestrates distinct outputs, including pathogen-specific transcriptional programs. Overall, our study demonstrates the independent control of immunity and longevity by a conserved regulatory protein.

## INTRODUCTION

Recent advances in aging research have led to a shift in focus from increasing lifespan to improving healthspan, a concept that encompasses measures of physiological and behavioral health, including stress resistance (Keith et al., 2014; Melov, 2016; Richardson et al., 2016), and acknowledges the equal importance of both good health and long life (Sierra, 2016). Lifespan, as a precise, measurable entity, has proven to be invaluable in classical genetic studies for identification of aging genes and pathways, including conserved factors that appear to promote longevity from worms to humans (Guarente and Kenyon, 2000; Martinez Corrales and Alic, 2020). A strong positive correlation between longevity and stress resistance has been observed, in model organisms and in nature, and many genes that increase lifespan have been reported to enhance stress resilience (Epel and Lithgow, 2014; Kirkwood et al., 2000; Murakami, 2006; Zhou et al., 2011), underscoring the relevance of lifespan to aging studies. However, in worms, flies and mice, descriptions of long-lived mutants that do not exhibit elevated stress resistance, and *vice versa*, have existed in literature for long, along with reports of pro-longevity genes that do not alter stress resistance (Arum and Johnson, 2007; Dues et al., 2017; Meissner et al., 2004). Recently, we and others have identified genes that promote longevity but repress stress resilience demonstrating that these attributes are physiologically distinct (Amrit et al., 2019; Kew et al., 2020). Nonetheless, a large fraction of known pro-longevity genes promote stress resistance (Zhou et al., 2011), but whether they govern these processes by shared or distinct mechanisms, and the underlying molecular details, remains largely unknown.

Immunity is central to an organism’s stress response and thus an integral aspect of healthspan (Keith et al., 2014; Richardson et al., 2016). Age-related increase in disease susceptibility occurs across species, including in humans (Nikolich-Zugich, 2018), as demonstrated by the ongoing COVID-19 pandemic’s disproportionate impact on the elderly (Cunha et al., 2020). While both the adaptive and innate immune systems undergo age-related changes, the latter is a major focus of contemporary aging biology because inflammaging, a derailment of the innate-immune system resulting in chronic inflammation, has been postulated to underlie age-related pathologies such as cardiometabolic disease, cancer and neurodegeneration (Franceschi and Campisi, 2014; Fulop et al., 2019; Fulop et al., 2018). Moreover, longevity-promoting genes and drugs such as FOXO3A and Metformin, respectively, have been shown to ameliorate the inflammaging profile of older immune cells towards that of a younger cohort (Bharath et al., 2020; Lee et al., 2013). To what extent such genes and drugs modulate immune status directly, and how this is connected to their longevity effects is unknown, and has topical relevance to human aging biology.

Studies in the nematode, *Caenorhabditis elegans*, have been instrumental in identifying fundamental aging mechanisms, including the discovery that signals from the reproductive system influence lifespan and stress resistance (reviewed in Amrit and Ghazi, 2017a). In worms, removal of a population of totipotent germline-stem cells (GSCs), precursors of all adult gametes, increases lifespan dramatically (Hsin and Kenyon, 1999). GSC-less animals also display extraordinary metabolic adaptability and resilience against stressors such as heat, DNA damage or infections by Gram-positive and Gram-negative pathogens (Alper et al., 2010; Amrit and Ghazi, 2017a). The increased longevity of germline-less worms is attributable to a network of transcription factors that is activated in the somatic cells when GSCs are ablated in the gonad, including DAF-16, worm homolog of FOXO3A (Amrit and Ghazi, 2017a; Ziv and Hu, 2011). Interestingly, many of these longevity-supporting transcription factors govern inducible, acute stress-response pathways such as heat-shock response (HSR, HSF-1), oxidative-stress response (OSR, SKN-1) and mitochondrial- and endoplasmic reticular-unfolded protein stress response (mtUPR and erUPR, ATFS-1 and XBP-1, respectively) (reviewed in Amrit and Ghazi, 2017a; Higuchi-Sanabria et al., 2018; Morimoto, 2020). Previously, we identified a group of nuclear hormone receptors (NHRs) critical for germline-less longevity, including NHR-49 (Ratnappan et al., 2014). *C. elegans* NHR-49 performs functions undertaken by Peroxisome Proliferator-Activated Receptor alpha (PPARα), a key regulator of energy vertebrate metabolism (Contreras et al., 2013; Kersten, 2014). PPARα agonists such as Fibrates are major hyperlipidemia drugs in humans that lower blood triglyceride levels making this gene attractive in the context of aging (Tenenbaum and Fisman, 2012). We showed that NHR-49 coordinately upregulated the expression of genes involved in mitochondrial fatty acid β-oxidation as well as lipid desaturation and elongation to preserve lipid homeostasis and promote longevity (Ratnappan et al., 2014). NHR-49 also orchestrates the transcriptional responses to acute starvation and oxidative stress (Goh et al., 2018; Van Gilst et al., 2005b). However, whether NHR-49 promotes the immunoresistance of GSC-less worms, and if it is widely deployed in the response against pathogens is unknown.

In worms, and in other organisms, many regulators of longevity and stress resistance have been shown to control these processes by cell non-autonomous mechanisms (reviewed in Medkour et al., 2016; Morimoto, 2020). For instance, the broadly expressed transcription factor DAF-16 is sufficient in intestinal cells to confer longevity on GSC-less worms (Libina et al., 2003), whereas, *Drosophila* dFOXO activity in the fat body increases lifespan (Hwangbo et al., 2004). Worm XBP-1 and mouse XBP-1s act in neuronal cells to modulate organismal erUPR and glucose metabolism, respectively (Taylor and Dillin, 2013; Williams et al., 2014). But, intestinal DAF-16 cannot confer heat resistance on GSC-less animals and provides little lifespan benefit to the insulin/IGF1 receptor, *daf-2*, mutants whose longevity is also completely DAF-16 reliant (Kenyon et al., 1993; Libina et al., 2003). Observations such as these suggest that regulatory factors may act in specific locations to modulate specific processes, or regulate the same process from different locations under different conditions. They underscore the equal importance of location and physiological context to gene function (Amelio and Melino, 2020; Ben-David and Amon, 2020; Uschner and Klipp, 2019). From this standpoint, site-of-action studies of proteins that influence lifespan, stress resistance and other health measures address the identities of tissues that are vital to lifespan vs. healthspan. NHR-49, similar to PPARα, is expressed in multiple tissues including the neurons, intestine, muscles and the epithelium-like hypodermis (Ratnappan et al., 2014). But, where the protein acts to modulate lifespan or any of the stress modalities under its control is unknown.

In this study, we describe a role for NHR-49 in the innate-immune response against the Gram-negative pathogen *Pseudomonas aeruginosa* in long-lived, germline-less animals as well as fertile adults. We demonstrate that NHR-49 is differentially influenced by GSC loss vs. pathogen exposure and promotes longevity and immunity by distinct mechanisms. While NHR-49 expression in any somatic tissue can rescue longevity, its control of immunity is more stringent and only neuron-derived protein could promote pathogen resistance in multiple genetic backgrounds. These distinct regulatory modes also extend to the expression of canonical NHR-49 target genes such as *acs-2* and *fmo-2*. We found neither target gene to be essential for resistance against *P. aeruginosa* infection, unlike their reported roles during infection by other pathogens. Hence, NHR-49 appears to establish distinct responses to short-term stimuli such as pathogen attack vs. long-term lifespan changes, and to organize pathogen-specific transcriptional programs.

## RESULTS

### NHR-49 Co-regulates the Expression of Genes Essential for Germline-less Longevity with DAF-16 and TCER-1

Previously, we demonstrated that *nhr-49* inactivation, either by RNAi or by mutation, completely abrogates the enhanced lifespan of *glp-1* mutants, a temperature sensitive, sterile model of GSC-less longevity (Arantes-Oliveira et al., 2003). To identify the transcriptional changes orchestrated by NHR-49 upon GSC removal, we compared the transcriptomes of *glp-1* vs. *nhr-49;glp-1* mutants using RNA-Seq. We found that NHR-49 controlled the transcriptional upregulation of 1120 genes (UP class) and downregulation of 1140 genes (DOWN class) in GSC-depleted worms (Fig. 1A, B, Table S1A, B). Since our previous studies had shown that *nhr-49* is transcriptionally up-regulated upon GSC removal by the joint action of two transcription factors, DAF-16 and TCER-1, (Ratnappan et al., 2014) we examined the overlap between NHR-49 UP and DOWN targets with genes whose expression is altered by DAF-16 and/or TCER-1, in *glp-1* mutants (Amrit et al., 2016). The overlap with genes specifically upregulated by either of these factors was significant, ranging from ∼14% (DAF-16-SPECIFIC UP) to ∼34% (TCER-1-SPECIFIC UP) (Fig. S1A). Strikingly, 53% of genes up-regulated in *glp-1* mutants by *both* DAF-16 and TCER-1 (JOINT UP) (Amrit et al., 2016) were also identified within the NHR-49 UP class (65/123, R factor 8.3, P <4.853e-45) (Fig. S1A). Notably, 35 of these NHR-49 UP genes have known functional roles in germline-less longevity as their RNAi inactivation was shown to shorten the lifespan of *glp-1* mutants in our previous studies (Table S2) (Amrit et al., 2016; Ratnappan et al., 2014). Amongst the genes down-regulated by DAF-16 and TCER-1 too, the shared group (JOINT DOWN) showed a much higher overlap with the NHR-49 DOWN class (∼36%, 26/73, R 5.5, P <2.094e-13) as compared to genes specifically down-regulated by either factor alone (Fig. S1B). Thus, NHR-49 targets share the largest overlap with genes whose expression is jointly regulated by DAF-16 and TCER-1. Additionally, our RNA-Seq analysis identified functionally relevant genes essential for the lifespan extension induced by germline loss.

**Fig. 1:**
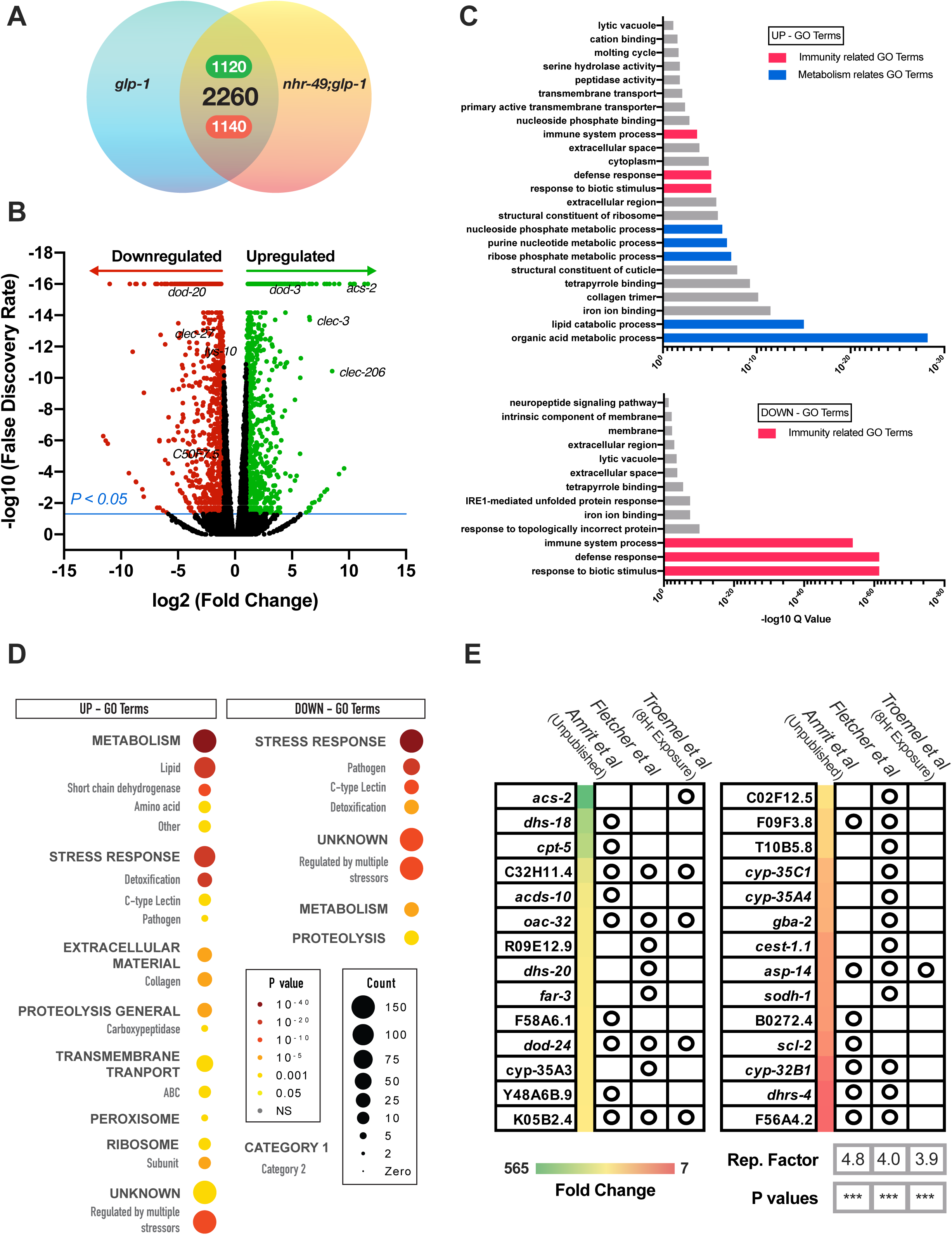
NHR-49 dictates a transcriptome upon germline loss that is enriched for innate-immunity genes. **(A)** Venn diagram depicting the genes upregulated (1120) and downregulated (1140) by NHR-49 following germline loss in *glp-1* mutants. **(B)** Volcano plot of gene expression changes between *glp-1* and *nhr-49;glp-1* animals. Differentially expressed genes highlighted as red (downregulated by NHR-49, DOWN) and green (upregulated by NHR-49, UP). Absolute log fold change >2, p value <0.05 and FDR of <2. **(C)** Wormbase Gene Set Enrichment Analysis (GSEA) of NHR-49 targets identified metabolism (blue) and stress response (red) categories as being enriched. **(D)** Gene ontology (GO) Term analysis using WormCat, a *C. elegans* identified pathogen response as one of the most enriched terms in both the UP class and DOWN Class. **(E)** Comparison of the top 100 UP NHR-49 targets with studies examining transcriptomic changes upon PA14 exposure {*Amrit et al., Unpublished* and (Fletcher et al., 2019; Shapira et al., 2006; Troemel et al., 2006; Twumasi-Boateng and Shapira, 2012)}.

### NHR-49 widely modulates the expression of innate-immunity genes

Based on NHR-49’s known role in lipid metabolism (Goh et al., 2018; Svensk et al., 2013; Van Gilst et al., 2005a), and our discoveries that the protein modulated the expression of fat metabolism genes in *glp-1* mutants (Ratnappan et al., 2014; Ratnappan et al., 2016), we anticipated NHR-49 targets to be enriched for lipid-metabolic functions. Gene Ontology (GO) analysis of the RNA-Seq data revealed that, while metabolic functions were indeed highly represented amongst NHR-49 targets, some of the most enriched GO terms related to stress response, especially immune response (Fig. 1C) (Amrit and Ghazi, 2017b). Analysis of this data through WormCat, a *C. elegans* bioinformatics platform that allows greater refinement of functional categories within enriched groups (Holdorf et al., 2020), substantiated these observations. Within the UP class, stress response, particularly pathogen response, was the second highest enriched category, and within the DOWN class, it was the most enriched one (Fig. 1D). Notably, 28 of the top 100 genes within the NHR-49 UP class were previously identified in studies examining gene-expression changes in worms infected by the human opportunistic pathogen *Pseudomonas aeruginosa* (Fig. 1E) {*Amrit et al., Unpublished* and (Fletcher et al., 2019; Shapira et al., 2006; Troemel et al., 2006; Twumasi-Boateng and Shapira, 2012)}. Together, these data suggested that besides lipid metabolism, NHR-49 also has a broad impact on expression of stress-response genes, especially genes involved in pathogen resistance. They motivated us to ask if NHR-49 had a functional role in *C. elegans* immunity.

### NHR-49 is essential for defense against *P. aeruginosa* pathogen attack

In order to address the role of NHR-49 in innate immunity, we tested the impact of *nhr-49* inactivation upon worm defense against *Pseudomonas*. We used survival following exposure to *Pseudomonas aeruginosa* strain PA14 (henceforth PA14) using the Slow Killing (SK) paradigm, wherein PA14 causes *C. elegans* to die over the course of several days (Keith et al., 2014; Powell and Ausubel, 2008). As previously reported, *glp-1* mutants survived significantly longer than wild-type (WT) adults (Alper et al., 2010; Evans et al., 2008). However, in *nhr-49;glp-1* mutants this resistance was abrogated. *nhr-49* single mutant^{\prime}s survival was significantly reduced compared to WT as well (Fig. 2A). An NHR-49 gain-of-function (gof) allele, *et7* (Lee et al., 2016; Svensk et al., 2013), produced a modest but inconsistent increase in survival in the presence of pathogen (Fig. S2A-C). In light of the functional homology between NHR-49 and mammalian PPARα, we asked if treatment with a PPARα agonist can improve worm immunity. Before and during infection with PA14, supplementation of worm food with Fenofibrate, a widely prescribed, lipid-lowering drug that acts by activating PPARα (Tenenbaum and Fisman, 2012), increased the survival of worms modestly (Figs. 2B and S2D, E). This improvement was abolished in the absence of *nhr-49* (Figs. 2B and S2D, E) suggesting that Fenofibrate acts through NHR-49 to augment immunoresistance. Hence, NHR-49 promotes longevity and immunoresistance in normal fertile *C. elegans* as well as germline-less, long-lived animals.

### Pathogen exposure causes reduction in NHR-49 protein, but not mRNA, levels

*nhr-49* mRNA and protein levels are both elevated in response to germline ablation (Ratnappan et al., 2014), so we asked how pathogen exposure impacted it. Using quantitative PCR (Q-PCR), we were unable to detect any change in *nhr-49* mRNA levels between worms fed OP50 vs. PA14 bacteria (Fig. 2C). Further, *nhr-49* was not identified as a gene whose expression was altered by PA14 exposure in five independent studies mapping PA14-induced transcriptomic changes (Fig. 1E) {*Amrit et al., Unpublished* and (Fletcher et al., 2019; Shapira et al., 2006; Troemel et al., 2006; Twumasi-Boateng and Shapira, 2012)}. We measured the levels of NHR-49 protein by using a transgenic strain expressing GFP-linked NHR-49 under control of its endogenous promoter. Both visual examination and automated quantification of fluorescence intensity using a COPAS^TM^ BIOSORT platform (Pulak, 2006) showed that GFP levels were significantly diminished in *glp-1* mutants exposed to PA14 (Fig. 2D-H). In fertile animals, PA14 infection induced a modest reduction that was visually evident but did not attain statistical significance (Fig. 2D-H). Together, these data suggest that unlike germline ablation that triggers both transcriptional and translational upregulation of NHR-49, infection with PA14 causes at least a modest reduction in protein levels.

**Fig 2.**
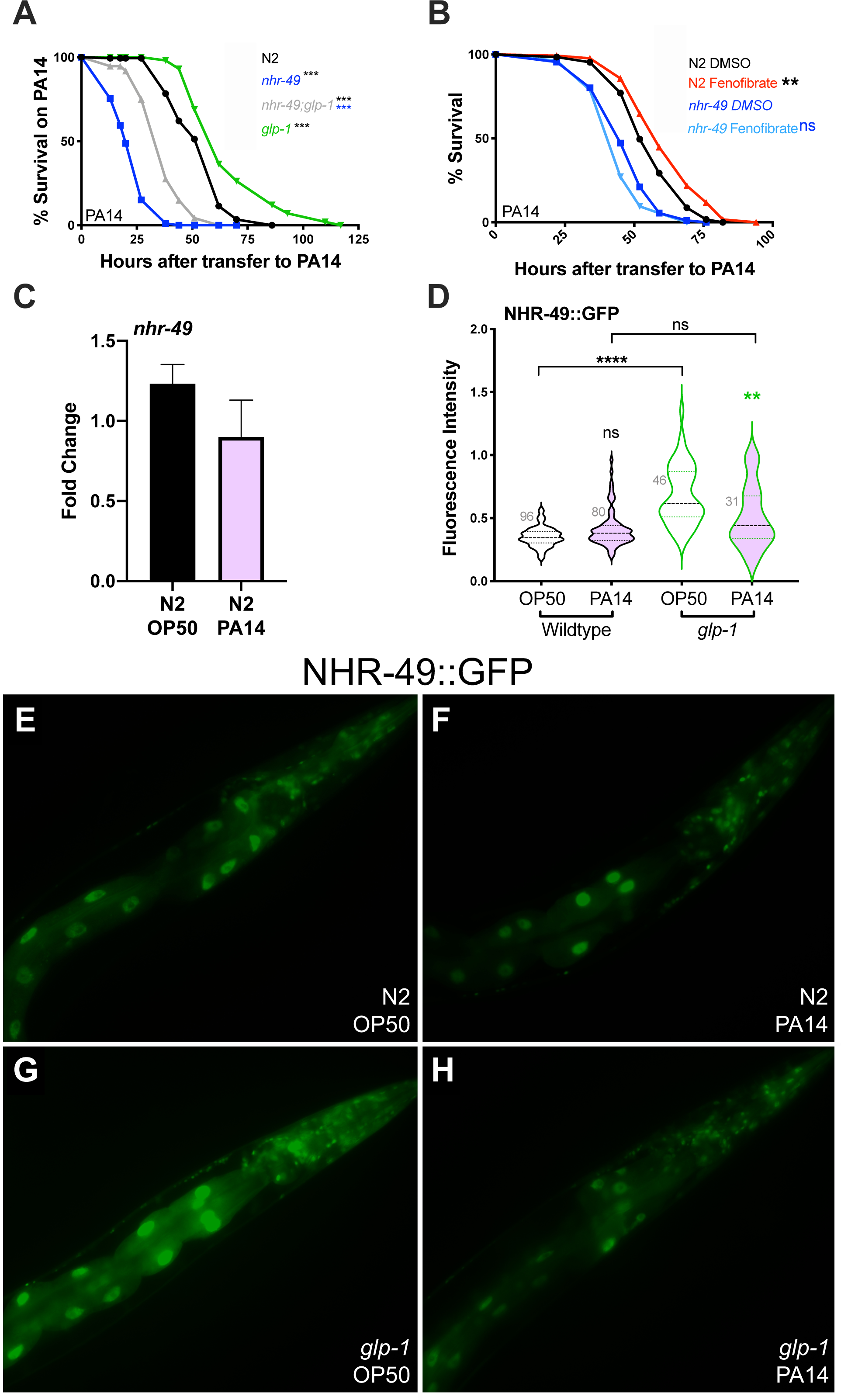
NHR-49 is essential for defense against *P. aeruginosa* pathogen attack. **(A)** Survival on PA14 of L4 stage wild-type worms (N2, black, m = 54.03 ± 0.99, n = 151/198) and well as *nhr-49* (blue, m = 23.03 ± 0.62, n = 184/195), *glp-1* (green, m = 67.31 ± 1.86, n = 101/113) and *nhr-49;glp-1* (gray, m = 37.48 ± 1.02, n = 99/115) mutants. See Tables S3 and S4 for additional trials with these strains. **(B) Fenofibrate increases immunity.** Survival on PA14 of L4 stage DMSO-control grown, wild-type worms (N2, black; m = 56.81 ± 1.08, n = 103/129) and *nhr-49* mutants (dark blue, m = 47.69 ± 0.98, n = 107/134) on PA14 compared to Fenofibrate-supplemented WT worms (red, m = 62.1 1.26, n = 93/135) and *nhr-49* mutants (light blue, m = 45.33 ± 0.89, n = 104/134). Data from additional trials in Fig. S2. (**C-H) PA14 exposure reduces NHR-49 levels. (C)** *nhr-49* mRNA levels measured by QPCR in Day 1 adults grown on OP50 till L4 stage then transferred to PA14 plates for 8 hours (pink) or continued on OP50 (black). P=0.27. Data combined from three independent biological replicates, each including three technical replicates. **(D)** Violin plot showing GFP intensity of NHR-49::GFP quantified on a COPAS Biosorter in Day 2 adults of wild-type animals (black outlines) and *glp-1* mutants (green outlines) grown on OP50 till L4 stage and then transferred to PA14 (pink) or retained on OP50 (blank) for 24h. Number of worms assayed shown on the panel. Data from one of three biological replicates with similar results. **(E-H):** Representative images from the experiment in C. Strains and conditions labeled on images. N2: wild type. In A and B, survival data shown as mean lifespan in hours (m) ± SEM (see methods for details). In C, error bars represent standard error of the mean (SEM). In D, where all fluorescence measures were normalized to time-of-flight, the center dashed line indicates mean intensity and flanking lines SEM. Statistical significance was calculated in A and B using the log rank method (Mantel Cox, OASIS2), in C by using a two-tailed t-test, and in D using a one-way non-parametric ANOVA with Dunn’s post hoc test (Graphpad Prism). Statistical significance is shown on each panel next to a given strain/condition with the color of the asterisk indicating strain/condition being compared to. P <0.01 (**), <0.001 (***), non-significant (ns).

### In germline-less animals, NHR-49 promotes immunity from neurons but longevity from multiple tissues

NHR-49 is widely expressed in *C. elegans* somatic cells in the intestine, muscles, hypodermis and neurons (Ratnappan et al., 2014). Previously, we found that NHR-49 expression under control of its endogenous/native promoter completely rescued the short lifespan of *nhr-49;glp-1* mutants to *glp-1* levels when animals were fed the normal diet of *Escherichia coli* strain OP50 (henceforth OP50). We asked if a similar widespread expression rescued the exceptionally short survival of *nhr-49;glp-1* mutants on a PA14 pathogenic diet as well. Surprisingly, endogenous-promoter driven NHR-49 not only failed to improve the survival of *nhr-49;glp-1* mutants on PA14, it reduced it even further (Fig. 3A, Table S3A). Animals carrying the same transgene showed consistent rescue of lifespan on OP50 (Fig. 3B, Table S3A). We checked if PA14 exposure abolished expression from the transgene explaining the lack of rescue. But, although intensity of the fluorescence signal in the GFP-tagged transgene was slightly reduced (as predicted by expression analysis described above), it was widely visible in all tissues. Additionally, lower NHR-49 levels did not explain the further aggravation in pathogen susceptibility we observed. This could not be attributed to transgene toxicity either because expression using the same transgene did not cause PA14 susceptibility when expressed in other genetic backgrounds (see sections below and Discussion).

**Fig. 3.**
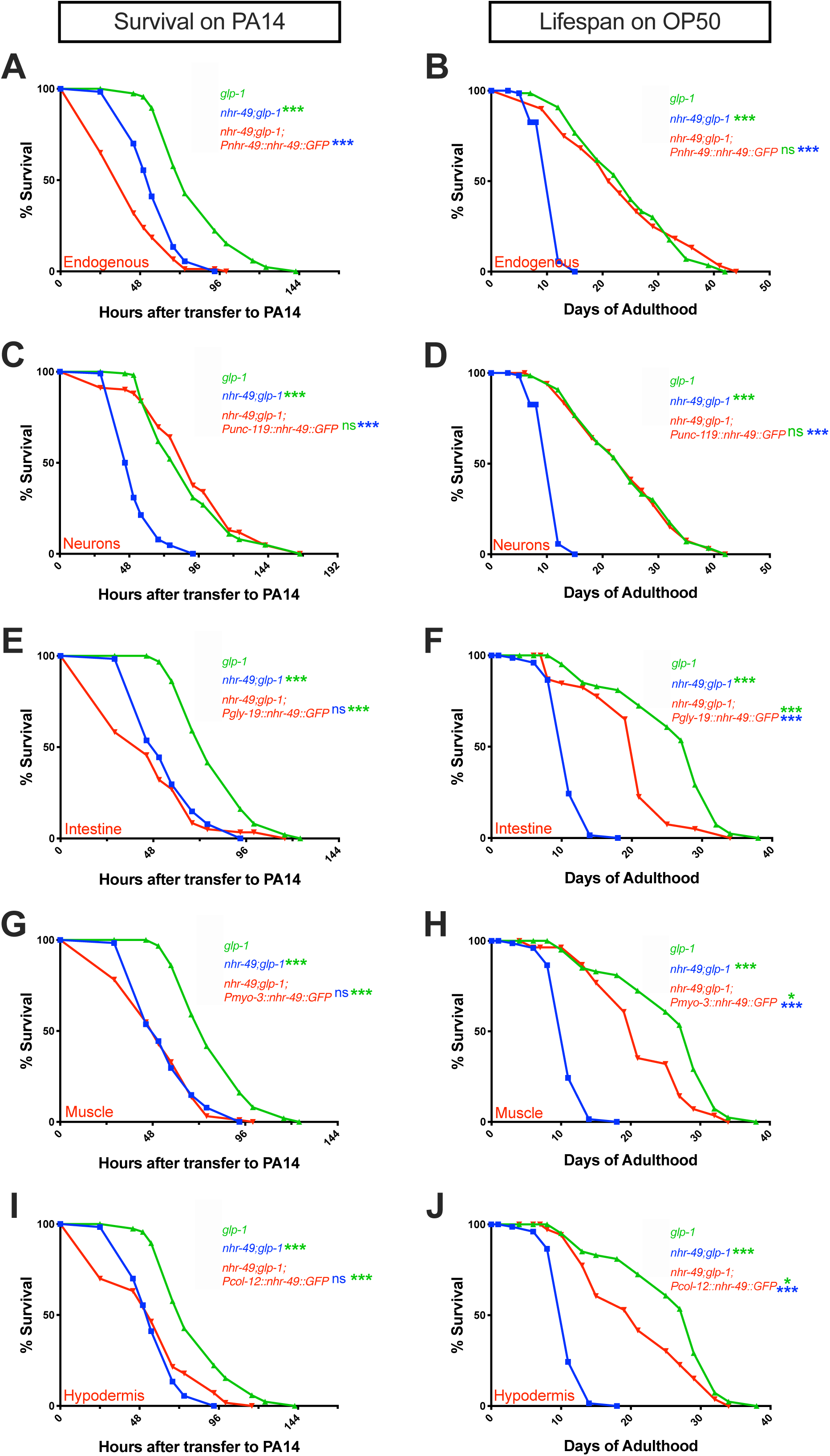
In germline-less animals, NHR-49 acts cell non-autonomously to promote immunity from neurons but longevity from multiple tissues. **(A, C, E, G, I) NHR-49 expression in neurons alone rescues PA14 resistance of *nhr-49;glp-1* mutants.** Mean PA14 survival (in hours) of *glp-1* (green), *nhr-49;glp-1* (blue), and *nhr-49;glp-1* mutants expressing NHR-49 in different tissues (red). **(A, I)** Survival of *glp-1* (82.88 ± 2.08, n= 105/120) and *nhr-49;glp-1* (57.92 ± 1.35, n= 108/120) mutants compared to transgenic *nhr-49;glp-1* mutants expressing NHR-49 via **Endogenous promoter** (**A**, 43.33 ± 1.74, n= 95/120) or in the **Hypodermis** (**I**, 56.06 ± 2.53, n= 96/120). **(C)** Survival of *glp-1* (89.65 ± 2.83, n= 104/108), *nhr-49;glp-1* (53.32 ± 1.26, n= 98/111), and *nhr-49;glp-1* mutants expressing NHR-49 in **Neurons** (**C**, 92.09 ± 3.49, n= 90/102). **(E, G)** Survival of *glp-1* (80.71 ± 1.62, n= 117/124), *nhr-49;glp-1* (56.25 ± 1.42, n= 111/121), and *nhr-49;glp-1* mutants expressing NHR-49 in **Intestine** (**E**, 47.82 ± 2.57, n= 68/79) and **Muscles** (**G**, 52.44 ± 1.67, n= 109/115). **(B, D, F, H, J) NHR-49 expression in any somatic tissue substantially rescues longevity of *nhr-49;glp-1* mutants on OP50.** Mean OP50 lifespan (in days) of *glp-1* (green), *nhr-49;glp-1* (blue) and *nhr-49;glp-1* mutants expressing NHR-49 in different tissues (red). **(B, D)** Lifespan of *glp-1* (24.43 ± 1.08, n= 51/70) and *nhr-49;glp-1* (11.28 ± 0.26, n= 55/72) mutants compared to transgenic *nhr-49;glp-1* mutants expressing NHR-49 via **Endogenous promoter** (**B**, 23.72 ± 1.32, n= 59/60) or promoters expressed in **Neurons** (**D**, 24.41 ± 1.04, n= 66/71). **(F, H, J)** Lifespan of *glp-1* (25.78 ± 0.98, n= 65/77), *nhr-49;glp-1* (11.27 ± 0.26, n= 72/72), and *nhr-49;glp-1* mutants expressing NHR-49 in **Intestine** (**F**, 19.79 ± 0.95, n= 41/63), **Muscles** (**H**, 21.4 ± 1.02, n= 34/59) or **Hypodermis** (**J**, 21.06 ± 1.36, n= 28/42). Survival and lifespan data shown as mean ± standard error of the mean (SEM). ‘n’ refers to number of worms analyzed over total number of worms tested in the experiment (see Methods for details). Statistical significance was calculated using the log rank (Mantel Cox) method and is indicated by asterisks on each panel next to mutant name (color of asterisk indicates strain being compared to). p<0.05 (*), <0.001 (***), not significant (ns). Note: In some panels (A, I; E, G; B, D; F, H, J), assays have the same controls are they were performed in the same biological replicate. Data from additional trials of these experiments (and wild-type controls) are presented in Table S3A-E.

The contradictory observations with the endogenous promoter could be explained if NHR-49’s impact on immunoresistance is tissue specific with expression in some sites exerting pro-immunity effects and in others reducing immunity. To test the possibility of tissue-specific functions, we expressed NHR-49 in individual tissues of *nhr-49;glp-1* mutants and examined the effect on their survival upon PA14 infection (Fig. 3C, E, G, I, Table S3B-E) as well as lifespan on a normal OP50 diet (Fig. 3D, F, H, J, Table S3B-E). We found that NHR-49 expression selectively in the neurons of *nhr-49;glp-1* mutants (using the *unc-119* promoter) (Maduro and Pilgrim, 1995) completely and reliably rescued their survival on PA14 to the same level as *glp-1* mutants (Fig. 3C, Table S3B). Expression in other somatic tissues had marginal and inconsistent impacts. Intestinal expression (*gly-19* promoter) (Warren et al., 2001) did not produce any significant increase in survival in any of 3 trials (Fig. 3E, Table S3C), whereas, hypodermal expression (*col-12* promoter) (Pujol et al., 2008) or expression in muscles (*myo-3* promoter) (Fire and Waterston, 1989) showed sporadic rescues (Fig. 3G, I, Table S3D, E). Thus, NHR-49 expression in neurons alone was sufficient to completely and reliably rescue the PA14 susceptibility of *nhr-49;glp-1* mutants.

We next asked how NHR-49 expression in individual tissues (using the same promoters as above) impacted *nhr-49;glp-1* mutant’s lifespan on a diet of OP50. Pan-neuronal expression completely rescued lifespan to *glp-1* levels in every trial (Fig. 3D, Table S3B). Interestingly, expression in each of the other three somatic tissues also produced substantial increases in longevity (Figs. 3F, H, J, Table S3C-E), although rescue to *glp-1* levels was achieved by neuronal NHR-49 alone. Together, these experiments showed that upon germline loss, while NHR-49 can act from any somatic tissue to largely promote longevity, only neuronal expression promotes PA14 resistance completely and reliably.

### In *nhr-49* mutants, hypodermal NHR-49 rescues longevity but diminishes immunity

Since *nhr-49* single mutants also exhibit shortened survival as compared to WT controls, we investigated which tissues NHR-49 acted in to promote their survival on PA14 and OP50. However, unlike in the *nhr-49;glp-1* background, the endogenous-promoter driven NHR-49 transgene rescued the survival of *nhr-49* single mutants on PA14 reliably (98% rescue in 3/4 trials, Fig. 4A, Table S4A) as well as on OP50. In fact, lifespan on OP50 was augmented even further as observed in our previous work (Fig. 4B, Table S4A) (Ratnappan et al., 2014). As in *nhr-49;glp-1* mutants, pan-neuronal expression completely rescued both the immunoresistance and lifespan of *nhr-49* mutants suggesting a key role for neurons in both immunity and longevity (Fig. 4C, D, Table S4B). Additionally, intestinal expression also significantly rescued both longevity and immunity (Fig. 4E, F, Table S4C), whereas, muscle expression rescued neither (Fig. 4G, H, Table S4D). Strikingly though, hypodermal NHR-49 completely rescued longevity on OP50, but survival on PA14 was significantly worsened (∼19% reduction in 4/4 trials, Fig. 4I, J, Table S4E). These results, along with the observations above, suggest that NHR-49 acts cell non-autonomously to modulate both longevity and immunity. While its function in multiple somatic tissues promotes longevity, the impact on immunoresistance is much more stringent, driven by neuronal activity and governed by the presence or absence of the germline.

**Fig 4.**
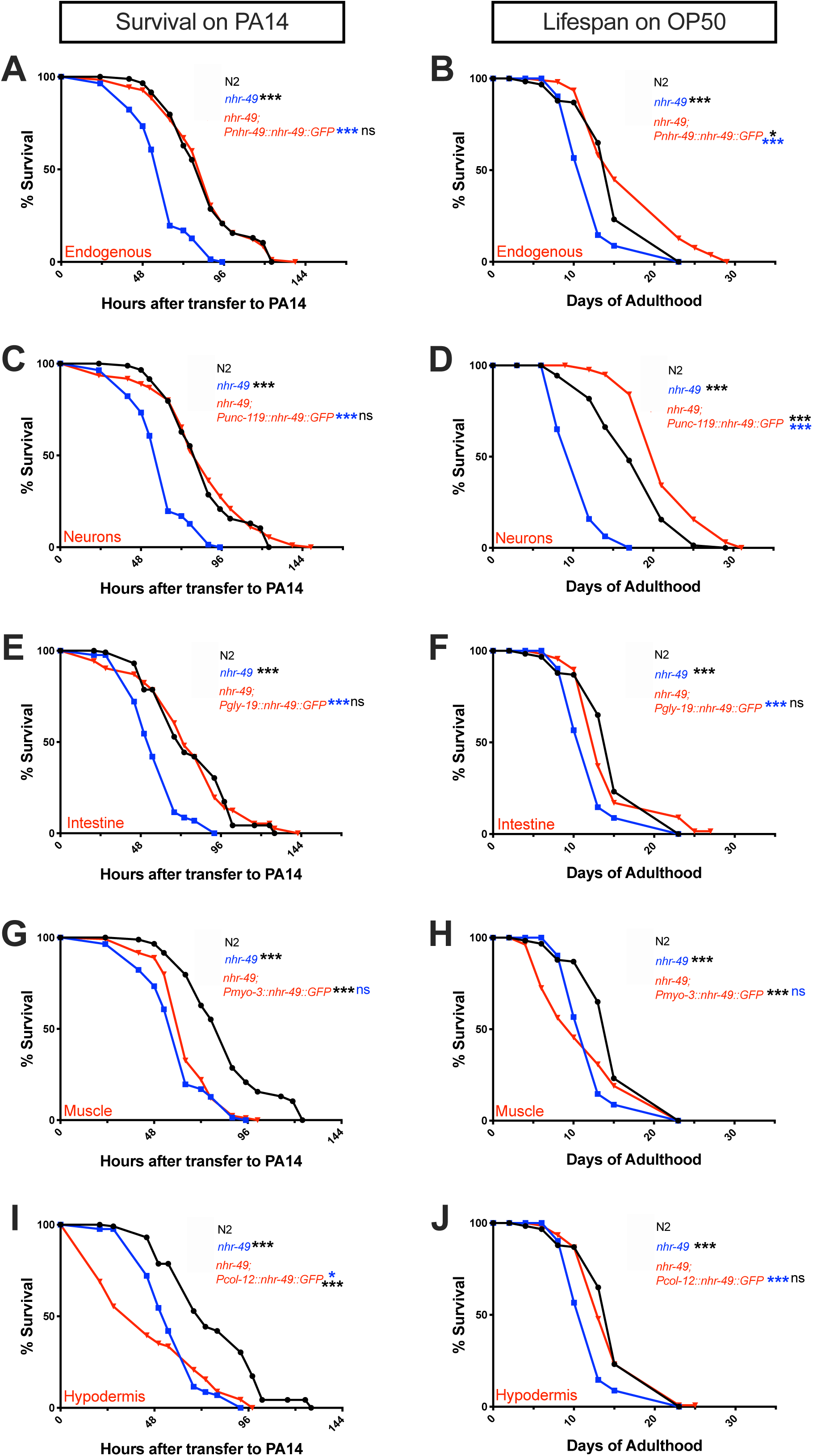
In *nhr-49* mutants, neuronal NHR-49 rescues both lifespan and immunity while hypodermal expression rescues longevity but lowers immunity. **(A, C, E, G, I) NHR-49 expression in neurons and intestine rescues the resistance of *nhr-49* mutants on PA14.** Mean PA14 survival (in hours) of WT (black, N2), *nhr-49* (blue), and *nhr-49* mutants expressing NHR-49 in different tissues (red). **(A, C, G)** Survival of N2 (84.82 ± 2.63, n= 55/90) and *nhr-49* (60.24 ± 1.52, n= 102/113) strains compared to *nhr-49* mutants expressing NHR-49 via **Endogenous promoter** (**A**, 84.35 ± 2.17, n= 100/125) or promoters expressed in the **Neurons** (**C**, 84.44 ± 2.73, n= 95/110) or **Muscles** (**G**, 65.73 ± 1.27, n= 105/119). **(E, I)** Survival of N2 (78.24 ± 2.57, n= 63/125), *nhr-49* (58 ± 1.44, n= 99/127), and *nhr-49* mutants expressing NHR-49 in **Intestine** (**E**, 77.6 ± 2.73, n = 87/125) or **Hypodermis** (**I**, 46.27 ± 2.72, n= 79/100). (B, D, F, H, J) NHR-49 expression in the neurons, intestine or hypodermis substantially improves *nhr-49* mutant’s longevity on OP50. Mean lifespans on OP50 (in days) of WT (black, N2), *nhr-49* (blue), and *nhr-49* strains expressing NHR-49 in different tissues (red). **(B, F, H, J)** Lifespan of N2 (15.42 ± 0.49, n= 89/122) and *nhr-49* (12.5 ± 0.36, n= 110/119) strains compared to *nhr-49* mutants expressing NHR-49 via **Endogenous promoter** (**B**, 18 ± 0.97, n= 97/112) or promoters expressed **Intestine** (**F**, 15.25 ± 0.4, n= 97/118), **Muscles** (**H**, 12.01 ± 0.71, n= 75/80) or **Hypodermis** (**J**, 14.86 ± 0.45, n= 131/141). **(D)** Lifespan of N2 (17.99 ± 0.58, n= 71/82) and *nhr-49* (11.11 ± 0.33, n= 63/65) strains compared to *nhr-49* mutants expressing NHR-49 in **Neurons** (**D**, 22.23 ± 0.71, n= 33/76). Survival and lifespan data shown as mean ± standard error of the mean (SEM). ‘n’ refers to number of worms analyzed over total number of worms tested in the experiment (see Methods for details). Statistical significance was calculated using the log rank (Mantel Cox) method and is indicated by asterisks on each panel next to mutant name (color of asterisk indicates strain being compared to). p<0.05 (*), <0.001 (***), not significant (ns). Note: assays in some panels (A, C, G; E, I; B, F, H, J) have the same controls are they were performed in the same biological replicate. Data from additional trials of these experiments and wild-type controls are presented in Table S4A-E.

### Elevating NHR-49 levels in neurons, or intestine, enhances immunoresistance but not longevity

NHR-49 protein levels are important in determining the animals’ lifespan because elevating its levels in normal, fertile adults induces a modest lifespan extension (Ratnappan et al., 2014). We asked if it increased PA14 resistance as well but found the answer to be equivocal because survival on PA14 was increased in 1/3 trials (Fig. 5A, B, Table S5A). We next asked if elevating NHR-49 levels in individual tissues could enhance immunoresistance (especially in the neurons that appeared to be the major site for its pro-immunity functions). Surprisingly, WT animals’ immunity was enhanced by NHR-49 overexpression in the neurons or the intestine (∼15-30%) (Fig. 5C, E, Table S5B, C, respectively). Muscle or hypodermal overexpression did not produce consistent changes in immunity (Fig. 5G, I, Table S5D, E). Interestingly, the benefits obtained by intestinal or neuronal upregulation were restricted to survival during PA14 infection. Lifespan on OP50 was not enhanced reliably by NHR-49 in any single somatic tissue (Fig. 5D, F, H, J, Table S5B-E). These results showed that elevating NHR-49 level in the precise location is important in its roles in immunity or longevity.

**Fig 5.**
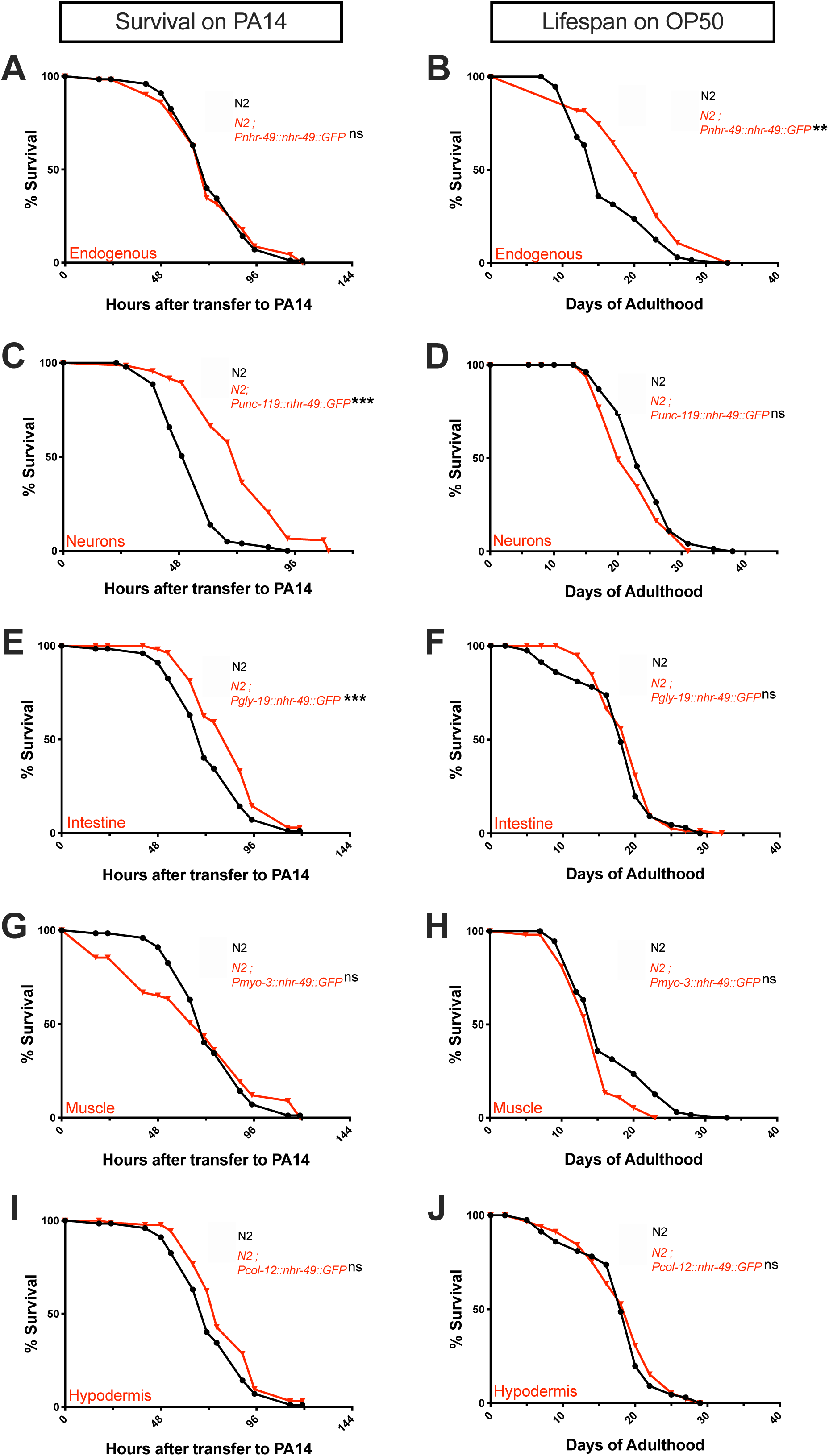
In wild-type animals, elevating NHR-49 levels in neurons or intestine enhances immunity but not longevity. **(A, C, E, G, I) NHR-49 overexpression in neurons or intestines increases immunity.** Mean survival on PA14 (in hours) of WT worms (black, N2) and strains overexpressing NHR-49 in different tissues (red). **(A, E, I, G)** Survival of N2 (76.98 ± 2.25, n= 102/129) and strains overexpressing NHR-49 via **Endogenous promoter** (**A**, 76.89 ± 3.06, n= 75/121) or promoters expressed in **Intestine** (**E**, 91.97 ± 2.65, n= 96/124), **Muscles** (**G**, 72.11 ± 4.28, n= 66/100) or **Hypodermis** (**I**, 82.68 ± 2.65, n= 63/97). **(C)** Survival of N2 (54.32 ± 1.09, n= 125/145) and strain overexpressing NHR-49 in **Neurons** (**C**, 73.72 ± 1.61, n= 114/136). **(B, D, F, H, J) NHR-49 upregulation in individual somatic tissues does not enhance immunity.** Mean lifespan on OP50 (in days) of WT worms (black, N2) and strains overexpressing NHR-49 in different tissues (red). **(B, H)** Lifespan of N2 (16.67 ± 0.65, n= 67/73) and strains overexpressing NHR-49 via **Endogenous promoter** (**B**, 20.83 ± 1.01, n= 35/44) or promoter expressed in **Muscles** (**H**, 14.6 ± 0.55, n= 39/52). **(D)** Lifespan of N2 (24.2 ± 0.55, n= 74/91) and strain overexpressing NHR-49 in **Neurons** (**D**, 22.35 ± 0.6, n= 57/94). **(F, J)** Lifespan of N2 (17.81 ± 0.56, n= 73/121) and strains overexpressing NHR-49 in **Intestine** (**F**, 19.06 ± 0.4, n= 86/116) or **Hypodermis** (**J**, 18.3 ± 0.53, n= 95/120). Survival and lifespan data shown as mean ± standard error of the mean (SEM). ‘n’ refers to number of worms analyzed over total number of worms tested in the experiment (see Methods for details). Statistical significance was calculated using the log rank (Mantel Cox) method and is indicated by asterisks on each panel next to mutant name (color of asterisk indicates strain being compared to). p<0.05 (*), <0.001 (***), not significant (ns). Note: assays in some panels (A, I, E, G; B, H; F, J) have the same controls are they were performed in the same biological replicate. Data from additional trials of these experiments and controls are presented in Table S5A-E.

Lastly, we assessed the consequences of raising NHR-49 levels in animals that have elevated protein to begin with, i.e., in *glp-1* mutants wherein NHR-49 is both transcriptionally and translationally upregulated (Ratnappan et al., 2014). Further overexpression in this genetic background, either using the widespread endogenous promoter or tissue-specific drivers, did not improve longevity or immunity. In fact, it appeared to diminish survival, especially upon PA14 exposure and in some cases on OP50 as well (Fig. 6A-J, Table S6A-E). Altogether, the above experiments substantiated the significance of both levels and location of NHR-49 in determining its ultimate impact on the lifespan and immune status of the animal. They suggested that a precise calibration of its tissue-specific expression is critical for its pro-immunity function.

**Fig 6.**
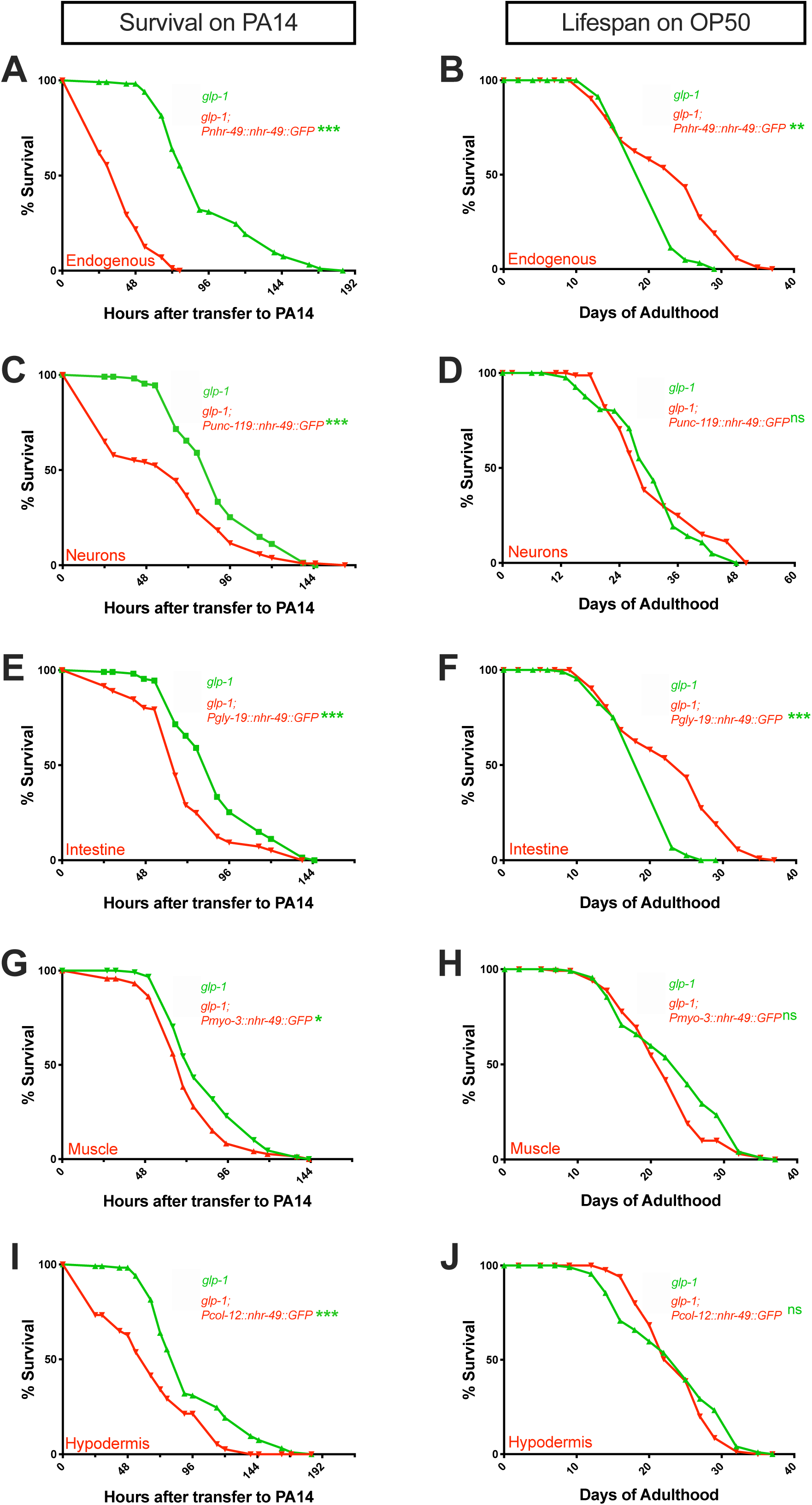
Upregulating NHR-49 in germline-less animals reduces immunity. **(A, C, E, G, I) NHR-49 overexpression in tissues of *glp-1* mutants reduces PA14 resistance.** Mean survival on PA14 (in hours) of *glp-1* (green) and *glp-1* mutants expressing NHR-49 in different tissues (red). **(A, I)** Survival of *glp-1* (94.91 ± 3.2, n= 102/117) compared *glp-1* expressing NHR-49 via **Endogenous promoter** (**A**, 39.35 ± 1.56, n= 94/105) or promoter expressed in the **Hypodermis** (**I**, 65.17 ± 4.78, n= 47/60). **(C, E)** Survival of *glp-1* (88.89 ± 2.57, n= 95/109) and *glp-1* expressing NHR-49 in **Neurons** (**C**, 59.86 ± 3.3, n= 111/120) or **Intestine** (**E**, 69.98 ± 2.52, n= 106/120). **(G)** Survival of *glp-1* (83.68 ± 2.04, n= 116/127) and *glp-1* expressing NHR-49 in **Muscles** (**G**, 73.05 2.12, n= 96/118). **(B, D, F, H, J) Effect of NHR-49 overexpression on *glp-1* mutant’s lifespan on OP50.** Mean lifespan on OP50 (in days) of *glp-1* (green) and *glp-1* mutants overexpressing NHR-49 in different tissues (red). **(B, F)** Lifespan of *glp-1* (22.98 ± 0.67, n= 122/124) compared to *glp-1* overexpressing NHR-49 via **Endogenous promoter** (**B**, 21.31 ± 0.42, n= 94/108) or promoter expressed in **Intestine** (**F**, 20.69 ± 0.44, n= 105/122). **(D)** Lifespan of *glp-1* (30.54 ± 0.79, n= 120/124) and *glp-1* overexpressing NHR-49 in **Neurons** (**D**, 31.4 ± 1.1, n= 70/80). **(H, J)** Lifespan of *glp-1* (23.35 ± 0.69, n= 101/114) and *glp-1* overexpressing NHR-49 in **Muscles** (**H**, 21.89 ± 0.55, n= 104/121) or **Hypodermis** (**J**, 23.79 ± 0.54, n=84/102). Survival and lifespan data shown as mean ± standard error of the mean (SEM). ‘n’ refers to number of worms analyzed over total number of worms tested in the experiment (see Methods for details). Statistical significance was calculated using the log rank (Mantel Cox) method and is indicated by asterisks on each panel next to mutant name (color of asterisk indicates strain being compared to). p<0.05 (*), <0.001 (***), not significant (ns). Note: assays in some panels (A, I; C, E; B, F; H, J) have the same controls are they were performed in the same biological replicate. Data from additional trials of these experiments and wild-type controls are presented in Table S6A-E.

### Expression of NHR-49-target genes, *fmo-2* and *acs-2*, is differentially impacted by germline loss and pathogen attack

We asked if the site-specific impacts of NHR-49 on longevity vs. immunity extended to its downstream targets too. Our NHR-49 UP group included the canonical *nhr-49*-target gene, *acs-2*, one of 22 *C. elegans* genes encoding Acyl CoA Synthetase (Van Gilst et al., 2005a; Van Gilst et al., 2005b), as well as *fmo-2*, that encodes a flavin monooxygenase (565 and 270 folds, respectively; Table S1A) (Goh et al., 2018; Leiser et al., 2015). In a recent study, both genes were reported to be dramatically upregulated upon infection by the Gram-positive pathogen, *Enterococcus faecalis*, and both were essential for survival during *E. faecalis* infection (Dasgupta et al., 2020). We asked if, and how, PA14 exposure altered their expression. *Pfmo-2p::GFP* was instead significantly downregulated upon PA14 exposure and independent of NHR-49 activity (Fig. 7A-E). *Pacs-2p::GFP* showed a small (<2 fold), NHR-49-dependent increase in expression in the presence of PA14 (Fig. 7F-J). Further, loss of function mutations in neither of the genes induced PA14 susceptibility; indeed *fmo-2* mutant showed a small but significant improvement in survival during infection (Fig. 7K) suggesting that NHR-49 exhibits specificity not only in the context of longevity vs. immunity, but also in a pathogen-specific manner.

**Fig 7.**
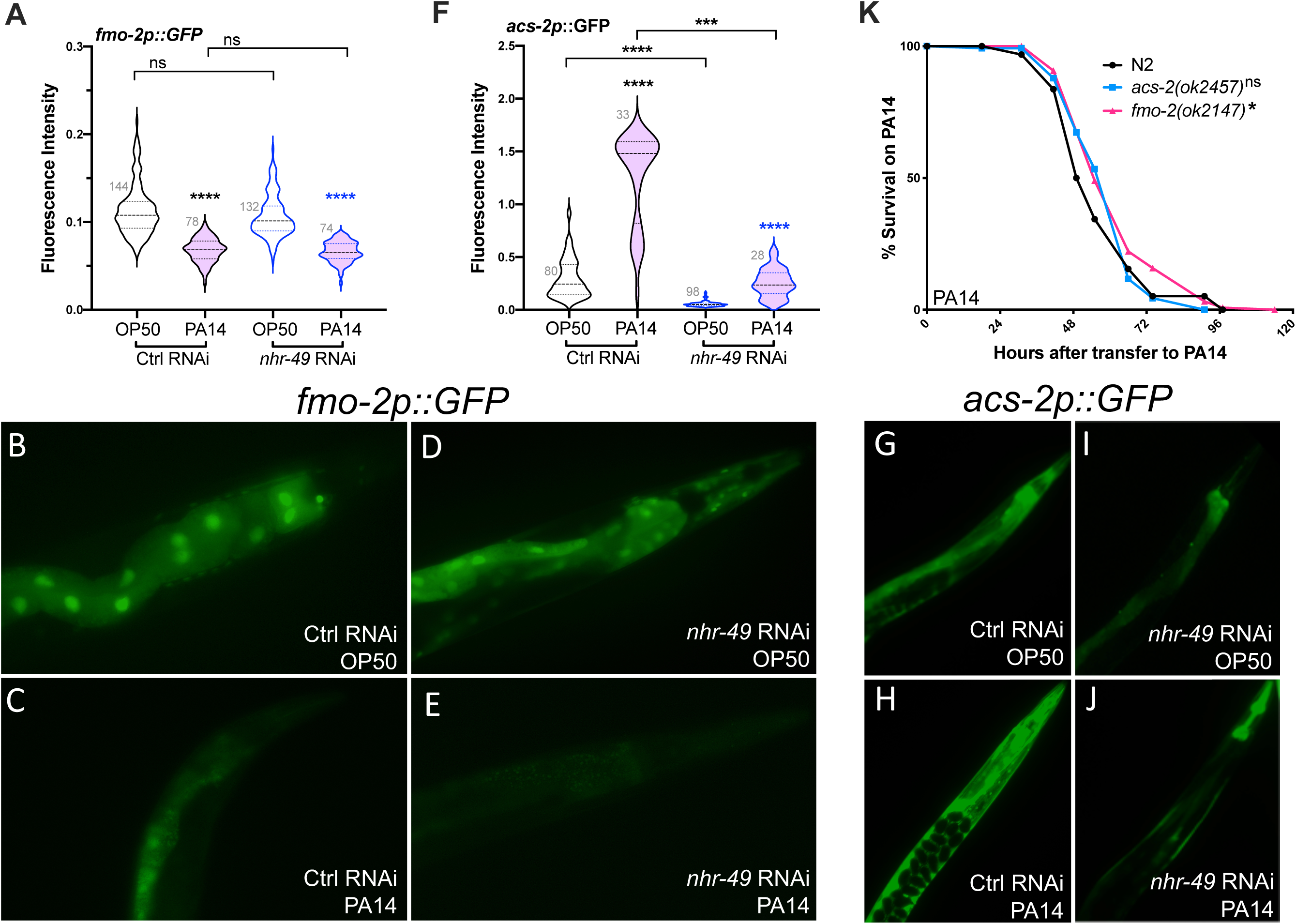
Expression of NHR-49-target genes, *fmo-2* and *acs-2*, is differentially impacted by germline loss and PA14 attack. **(A-E) *fmo-2* expression decreases upon PA14 exposure in an *nhr-49*-independent manner. (A)** Violin plot showing GFP intensity quantified on a COPAS Biosorter of Day 2 adult P*fmo-2p*::GFP animals grown to the Day 1 stage on vector control (black outline) or *nhr-49* RNAi (blue outline) bacteria before transfer to PA14 (pink) or OP50 (blank) for 24h. Number of worms assayed per condition shown. Data from one of three trials that gave similar results. **(B-E):** Representative images from experiment in A. Strains and conditions labeled on images. **(F-J) *acs-2* expression increases upon PA14 exposure in an *nhr-49*-dependent manner. (F)** Violin plot showing GFP intensity quantified on a COPAS Biosorter of Day 2 adult P*acs-2p*::GFP animals grown to the Day 1 stage on vector control (black outline) or *nhr-49* RNAi (blue outline) bacteria before transfer to PA14 (pink) or OP50 (blank) for 24h. Number of worms assayed per condition shown. Data from one of three trials that gave similar results. **(G-J):** Representative images from the experiment in F. Strains and conditions labeled on images. **(K)** Survival of L4 stage wild-type worms (N2, black) and *acs-2* (light blue), and *fmo-2* (fushia) mutants exposed to PA14. N2 (m = 56.66 ± 1.4, n = 97/127), *acs-2* (m = 59.52 ± 1.19, n= 94/137), and *fmo-2* (m = 62.53 ± 1.36, n = 136/153). In A and F, the center dashed line indicates mean intensity and lines flanking it represent standard error. In K, survival data shown as mean lifespan in hours (m) ± SEM (see methods for details). Statistical significance was calculated in A and F using a one-way non-parametric ANOVA with Dunn’s post hoc test (Graphpad Prism). Significance was calculated in K using the log rank method (Mantel Cox, OASIS2). Statistical significance shown on each panel with the color of the asterisk indicating the strain being compared to. P <0.01 (**), <0.001 (***), <0.0001 (****), non-significant (ns).

## DISCUSSION

In this study, we demonstrate that NHR-49 is a pro-longevity factor that modulates lifespan and immunity through distinct mechanisms. NHR-49 is influenced differently by, and orchestrates distinct responses to, the acute stress of pathogen attack vs. a lifespan-altering intervention such as germline loss. In particular, our site-of-action studies reveal the crucial significance of both location and physiological context in determining how regulatory proteins modulate individual measures of health and longevity.

### In Biology, Context is Critical

In the last decade, several studies have established the cell non-autonomous regulation of longevity and stress response, and identified tissues where key transcription factors act to modulate these processes (reviewed in Dillin et al., 2014; Medkour et al., 2016; Morimoto, 2020). But, the fact that in addition to site-of-expression, physiological context has a crucial role in determining whether a protein has beneficial, benign or detrimental impacts on longevity and immunity is poorly appreciated. A regulatory factor may act in different tissues to modulate different biological processes (Kodani and Nakae, 2020). Or, within the same tissue, the activity of a protein may have diametrically opposite impacts on different aspects of health (Amrit et al., 2019). Our experiments reveal this to be the case and show that even though NHR-49 enhances both longevity and immunity, it plays nuanced and complex roles in different somatic tissues in mediating its effects on these processes.

We found that NHR-49 expression in neurons alone could rescue the immunity deficits of germline-less, *nhr-49;glp-1* mutants, whereas, their lifespan was substantially restored by expression in any somatic tissue. In fertile, *nhr-49* single mutants, immunity was restored by presence in neurons or intestine, but lifespan could be rescued from other tissues as well. This suggests that pathogen response may be more sensitive to NHR-49’s location as compared to longevity. Interestingly, NHR-49 expression in muscles provided little or no immunity benefit in any of the genetic backgrounds we tested, irrespective of the presence or absence of the germline in the animal. In fact, expression in muscles mostly diminished immunity. This is in noticeable contrast to the broad immunity advantages conferred by neuronal NHR-49 in multiple genetic backgrounds. Of note, similar observations have been made with the erUPR regulator, XBP1, whose expression in neurons or intestine increases lifespan and proteostasis but expression in muscles diminishes both (Imanikia et al., 2019). Another intriguing observation in our study is the impact of widespread, endogenous promoter-driven NHR-49 expression on the PA14 resistance of *nhr-49;glp-1* mutants. Not only was this transgene unable to rescue the mutants’ PA14 sensitivity, it further diminished their survival upon infection. While it is possible that this is simply a consequence of transgene toxicity, it is unlikely because it is a functional transgene that (a) completely rescued the PA14 resistance of *nhr-49* single mutants (b) completely rescued the lifespan on OP50 of both *nhr-49;glp-1* and *nhr-49* mutants (c) did not cause lifespan shortening in normal animals fed either OP50 or PA14, and in fact extended lifespan on OP50. These data further emphasize that NHR-49’s site-and level-of expression are exquisitely sensitive during pathogen infection, and modest changes in either can have major consequences for the animal’s immune status.

Both intestine and hypodermis serve as fat depots in worms (Ashrafi, 2007; Watts and Ristow, 2017), so it was interesting to discover that the immunodeficiency of *nhr-49* mutants was rescued by intestinal NHR-49 expression but worsened further upon hypodermal expression. Recent work has described qualitative differences between the fats in these two sites. Intestinal fat is protein rich and largely destined for egg deposition, whereas hypodermal fat is less protein-encumbered and likely functions as the storage site (Chen et al., 2020). Considering the role of NHR-49 in lipid hydrolysis, it is enticing to speculate that it’s expression in these two tissues propels the mobilization of different lipid pools and consequent production of lipid signals that impact production of immune effectors differently.

### Pathogen-Specific Activity of NHR-49

Our data show that NHR-49 is differentially regulated by short-term pathogenic stress vs. lifespan altering intervention of germline loss, and in turn, it performs specific, tissue-dependent activities. A recent report identified *nhr-49* as being essential for defense against the Gram-positive pathogen, *E. faecalis* (Dasgupta et al., 2020), and our collaborator’s laboratory has identified a role for the gene in immunity against another Gram-positive bacterium, *Staphylococcus aureus* (*K Wani and J Irazoqui, Personal Communication, June 2020*) hinting at a broad role for the gene in innate immunity. Collectively though, evidence suggest that NHR-49 orchestrates pathogen specific transcriptional and functional outputs. For instance, the expression of canonical NHR-49 target genes, *acs-2* and *fmo-2*, is not only differently affected by germline loss vs. PA14 exposure, it is also altered in a pathogen-specific manner. Upon infection by *S. aureus* or *E. faecalis*, *fmo-2* expression is dramatically upregulated (>1000 fold and ∼150 fold, respectively) and *E. faecalis* exposure upregulates *acs-2* >1000 fold, in an *nhr-49*-dependent manner (Dasgupta et al., 2020) (*K Wani and J Irazoqui, Personal Communication, June 2020*). But, we found *fmo-2* to be downregulated by PA14 infection, whereas, *acs-2* showed a small (<2 fold) upregulation. Both genes are critical for survival upon *E. faecalis* exposure (Dasgupta et al., 2020), and *fmo-2* mutants are hyper-susceptible to *S. aureus* infection as well (*K Wani and J Irazoqui, Personal Communication, June 2020*). But, we found neither one contributed towards PA14 resistance and *fmo-2* mutants in fact survived longer than WT. These observations suggest that NHR-49 organizes a pathogen-specific, transcriptional program that determines the survival of the animal under different conditions. They indicate a molecular adaptability that can be highly advantageous to the animal in the ever-changing conditions in the wild.

How does NHR-49 activity in different somatic tissues produce such distinct consequences on immunity and longevity? It is likely through contextual expression of distinct sets of target genes in each of these tissues. For instance, DAF-16 has been shown to regulate the expression of targets in neurons that are distinct from those in other tissues such as the intestine (Kaletsky et al., 2016). The neuronal DAF-16 transcriptome includes genes required for memory and axon regeneration, whereas, the non-neuronal transcriptome includes longevity and metabolism genes. DAF-16 also appears to rely on different co-factors to mediate these tissue-specific gene expression changes, as its intestinal transcriptome is largely shared with PQM-1 (Tepper et al., 2013), whereas, neuronal genes are regulated by FKH-9 (Kaletsky et al., 2016). To understand NHR-49’s mode of action further, it is exigent to map the transcriptomes dictated by it in individual somatic tissues, in healthy as well as infected animals, and to identify transcriptional co-regulators it may be acting with in specific contexts. NHR-49 has been reported to partner with NHR-66 to regulate sphingolipid metabolism, and with NHR-80 to regulate fatty-acid desaturation (Pathare et al., 2012). We identified a group of 12 other NHRs that also promote germline-less longevity, and NHR-49 physically interacts with one of these, NHR-71, to control the expression of longevity genes (Ratnappan et al., 2016). These are attractive candidates for identifying tissue-specific and pathogen-specific NHR-49 transcriptional co-regulators in future studies.

### The Conservation of NHR-49’s Pro-Immunity Function

PPARα agonists such as Fibrates have been reported to increase lifespan and mitigate Amyloid β (Aβ)-induced paralysis in worms in an NHR-49-dependent manner (Leiteritz et al., 2020), strengthening the premise that the proteins share functional homology. Our observation here, on the role of NHR-49 in promoting immunity and its tissue-specific functions, are noteworthy in light of this and other similarities it shares with mammalian PPARα. Like NHR-49, PPARα plays well-defined roles in starvation-induced fatty-acid oxidation, oxidative stress, heat-stress resistance, lipoprotein metabolism and inflammatory responses (Bougarne et al., 2018; Christofides et al., 2020). Recently, intestinal PPARα signaling has been shown to be critical for maintaining immune tolerance towards commensal gut microbiota in mice (Manoharan et al., 2016). PPARα is also expressed in multiple tissues including the liver, brain, heart, muscles and immune cells (e.g., macrophages, monocytes, and lymphocytes). Importantly, PPARα performs distinct functions in these tissues and its activity is governed by tissue-specific mechanisms. For instance, it mediates fatty acid oxidation in liver and heart, but its endogenous ligands and their sources are different in the two organs. In liver, a lipid species, 16:0/18:1-GPC, derived by Fatty Acid Synthase (FAS)-mediated *de novo* lipogenesis, serves as its endogenous ligand (Chakravarthy et al., 2009), whereas, in the heart, triglyceride lipolysis by adipose triglyceride lipase (ATGL) generates ligands for PPARα activation (Haemmerle et al., 2011). Conversely, three endogenous PPARα lipid ligands have been identified in the brain where it regulates synaptic function and hippocampal plasticity (Roy et al., 2016). Our observation, that Fenofibrates enhance immunity against *P. aeruginosa*, is especially noteworthy and promising. Since NHR-49 appears to play a role in multiple bacterial infections, it will be interesting to ask if Fibrates increase resistance against other pathogens too, and if the immunity-promoting function of NHR-49 is shared with PPARα. Given the clinical success of targeting PPARα for anti-dyslipidemia drugs (Tenenbaum and Fisman, 2012), there is opportunity to assess Fenofibrates for potential immunity boosting functions and for intervention against inflammaging (Nogueira-Recalde et al., 2019).

## MATERIALS AND METHODS

### *C. elegans* strains and lifespan assays

All strains were maintained by standard techniques at 20°C or 15°C on nematode growth medium (NGM) plates seeded with an *E. coli* strain OP50. For experiments involving RNAi, NGM plates were supplemented with 1 mL 100 mg/mL Ampicillin and 1 mL 1M IPTG (Isopropyl β-D-1-thiogalactopyranoside) per liter of NGM. The main strains used in this study include N2 (wild type), CF1903 *[glp-1(e2144) III]*, AGP12a *[nhr-49(nr2041)I]*, AGP22 [*nhr-49(nr2041)I;glp-1(e2141)III]*, AGP110 *[nhr-49(et7)I].* All strains listed in Table S7. Lifespan experiments were performed as previously described (Amrit et al., 2014). For lifespan in the *glp-1* background, eggs were kept at 20°C for 2-6 h, grown to the L4 stage at 25°C, then shifted back to 20°C for remaining lifespan. In lifespan assays, the L4 stage was counted as Day 0 of adulthood. Fertile strains were transferred to fresh plates every other day to separate parents from progeny. Animals that exploded, bagged, crawled off the plate, or became contaminated were marked as censored upon observation. The program Online Application of Survival Analysis 2 (OASIS 2) (Han et al., 2016) was used for statistical analysis of both lifespan and pathogen stress assays. P-values were calculated using the log-rank (Mantel–Cox) test (Han et al., 2016). Results were graphed using GraphPad Prism (Version 8).

### Pathogen survival assays

Survival in the presence of the Gram-negative pathogen *Pseudomonas aeruginosa* strain PA14 was used in this study to assess the immunoresistance of *C. elegans* strains. Luria Bertani (LB) agar plates were streaked with bacteria from −80°C glycerol stocks, incubated at 37°C overnight and stored at 4°C for ≤ 1 week. Single PA14 colonies from streaked plates were then inoculated into 3mL King’s Broth (Sigma) overnight (16-18h) in a 37°C shaking incubator. 20 μL of this culture was seeded onto slow-killing (NGM with 0.35% peptone) plates and incubated at 37 °C for 24h (Keith et al., 2014; Tan and Ausubel, 2000). Seeded PA14 plates were kept at room temperature for 24h before use.

PA14 survival assays were performed as previously described (Keith et al., 2014). Age-matched *C. elegans* strains were grown under the same conditions and selected at the L4 stage as for the lifespan experiments. 25-30 L4 worms each were transferred to five PA14 plates per strain and maintained at 25°C till the end of their lives. Strains were monitored at 6-12h intervals to count living, dead, and censored animals as described above. Living animals were transferred to fresh PA14 plates each day for 3-4 days. Statistical analysis of survival data was performed on OASIS 2 and representative trials were graphed with GraphPad Prism (Version 8).

### RNA-Sequencing and data analysis

RNA was isolated from 3 biological replicates of Day 2 adults of CF1903 (*glp-1)* and AGP22 (*nhr-49;glp-1)* strains, grown as described above. Following 7 freeze thaw cycles, approximately 3000 worms were harvested for RNA using the Trizol method. RNA was checked for quality and quantity using the Agilent Tapestation and Qubit Fluorometry. Sequencing libraries were prepared using the TruSeq stranded mRNA (PolyA+) kit and the samples were then subjected to 75 base pair paired-end sequencing on an Illumina NextSeq 500 sequencer at the Univ. of Pittsburgh Genomics Research Core. Sequencing data was analyzed using the CLC Genomics Workbench (Version 20.0.3) employing the RNA Seq pipeline. Differentially regulated genes were filtered for significant changes based on the criteria of >2 fold change in expression, P Value of <0.05 and a false discovery rate (FDR) of <0.05.

### Gene Ontology analyses

Genes that were differentially regulated in a statistically significant manner were classified into two groups as either up-regulated (UP) or down-regulated (DOWN) NHR-49 targets. These groups were analyzed for enrichment of gene classes based on Gene Ontology (GO) Terms using *C. elegans* centered publicly available online resources, Wormbase Gene Set Enrichment Analysis tool (https://wormbase.org/tools/enrichment/tea/tea.cgi) and WormCat (http://wormcat.com/) (Holdorf et al., 2020). Representation Factor was calculated at http://nemates.org/MA/progs/overlap_stats.html.

### Q-PCRs

RNA was isolated as mentioned above, quantified using a Nanodrop and DNAse treated (DNAse kit, Sigma AMPD1). The RNA was then reverse transcribed into cDNA using the High-Capacity cDNA Reverse Transcription kit (Applied Biosystems, 4368814) following the manufacturer’s recommendations. RNA samples were collected from three independently isolated ‘biological replicates’, and three ‘technical replicates’ of each strain/condition were tested for a given biological replicate. Quantitative PCRs were conducted using the PowerUp SYBR Green Master Mix kit (Applied Biosystems A25741) on the CFX Connect Machine (BioRad). Gene expression data was analyzed using the ΔΔCt method and normalized to the housekeeping gene, *rpl-32*. Melt curves for all reactions were run confirming the integrity of the reaction. Primer sequences used for the experiment were *nhr-49_Fwd* (TTGGCAGAGGTGGATTCTC), *nhr-49_Rev* (CTGTAAAGAGACCGGAGCC), *rpl-32_Fwd* (GATTCCCTTGCGGCTCTT), and *rpl-32_Rev* (GATTCCCTTGCGGCTCTT).

### Transgenic strain generation

Tissue-specific NHR-49 expressing strains were generated by plasmid microinjection. A control P*nhr-49::nhr-49::gfp* (pAG4) construct, which drives NHR-49 expression via its endogenous promoter, was created as previously described (Ratnappan et al., 2014). To drive NHR-49 expression in other tissues, 4.4kb coding region of *nhr-49* was first amplified with modified primers to introduce SbfI and SalI restriction sites at the 5’ end and SmaI at the 3’ end of the coding region. This product was cloned into the GFP expression vector pPD95.77 (Addgene plasmid 1495) upstream of, and in frame with, GFP. Individual tissue-specific promoters were then amplified and ligated independently into this plasmid using primers modified with the forward primer including SbfI and the reverse including SalI to create plasmids for expressing NHR-49 in the muscle (P*myo-3*::NHR-49::GFP), intestine (P*gly-19*::NHR-49::GFP), hypodermis (P*col-12*::NHR-49::GFP) and neurons (P*unc-119*::NHR-49::GFP). Each of the five constructs were then injected at a concentration of 100 ng/mL with 15 ng/mL of P*myo-2::mCherry* co-injection marker into *nhr-49*, *nhr-49;glp-1, glp-1* and WT strains. For each of the 20 transgenic strains, 2-4 independent transgenic lines were generated. Transgene-carrying strains were maintained and selected for lifespan and pathogen stress assays using a Leica M165C microscope with a fluorescence attachment. A complete listing of all strains created for this study is provided in Table S7.

### Fenofibrate supplementation assay

100 μL of 10 μM Fenofibrate (Sigma F6020) in 0.1% DMSO were placed onto both NGM and slow-killing plates before seeding with OP50 or PA14, respectively, as described above (Brandstädt et al., 2013; Leiteritz et al., 2020). Upon the drying and growth of the bacterial lawn, eggs were grown to L4 on either the Fenofibrate or 0.1% DMSO control plates, then transferred to PA14 plates (similarly supplemented with Fibrate or DMSO) at L4 larval stage and survival monitored. Worms were transferred to fresh plates as described above.

### GFP fluorescence imaging and quantitation

GFP expression in transgenic strains was quantified using the COPAS Biosorter (Union Biometrica; Holliston) as described in (Pujol et al., 2008). For setup, progeny of transgenic mothers were grown to the Day 1 of adulthood (when fluorescent signal became clear) under normal conditions on *E. coli* OP50 or HT115 RNAi control strains at 20°C. For strains with the *glp-1* mutation, eggs were kept at 20°C for 2-6 h then grown to Day 1 at 25°C. Day 1 adults were transferred to PA14 plates or OP50 control plates and maintained 25°C for 24hrs before imaging and quantification. Each strain was washed into the COPAS sample cup with ∼5 mL deionized water for measurement of individual worms. Intensity of green fluorescence of each animal was normalized to the axial length measured (i.e. GFP fluorescence divided by time-of-flight). Statistical significance was determined using a one-way non-parametric ANOVA with Dunn’s post hoc test (Graphpad Prism). Representative whole-body images of transgenic worms were taken using 10 mM Sodium Azide for immobilization and imaged at 20x magnification using a Leica DM5500B compound scope with LAS X software (Leica).

## Acknowledgements

The authors are grateful to Javier Irazoqui (University of Massachusetts Medical School, Worcester) for sharing strains, reagents and unpublished data during this study. Some strains were provided by the CGC, which is funded by NIH Office of Research Infrastructure Programs (P40 OD010440). This work was supported by a grant from the National Institutes of Health (R01AG051659) to AG and a Children’s Hospital of Pittsburgh Research Advisory Committee (RAC) fellowship to NN.

## Author Contributions

AG conceived the project and designed the experiments; NN, FG, RR, JL, NB and AG performed the experiments; AG and NN wrote the manuscript with input from the other authors.

---

**Supplementary Figure 1:**
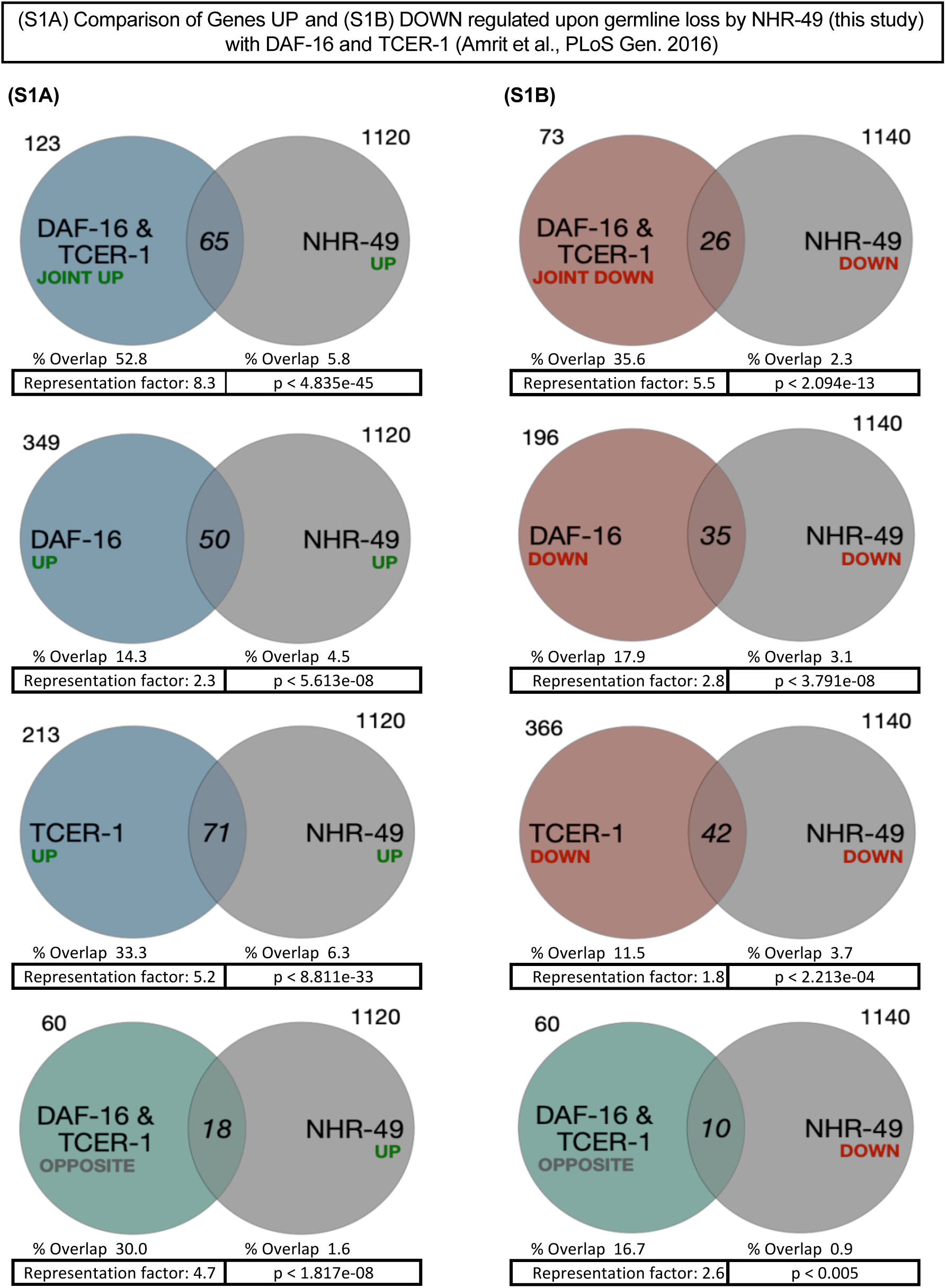
Overlap between NHR-49 target genes differentially expressed in *glp-1* mutants in this study with previously identified downstream targets of DAF-16 and TCER-1 in the same background (Amrit et al., 2016). **(A)** Comparison of genes upregulated by NHR-49 (UP) with genes upregulated by DAF-16 and TCER-1 jointly (Joint Upregulated: 53% Overlap, p < 4.835e-45) or by DAF-16 alone (Specific Upregulated: 14%, p < 5.613e-08), TCER-1 alone (Specific Upregulated: 33%, p < 8.811e-33) or upregulated by NHR-49 but oppositely impacted by DAF-16 vs. TCER-1 (Oppositely regulated: 30%, p < 1.817e-08). **(B)** Comparison of genes downregulated by NHR-49 (DOWN) with genes downregulated by DAF-16 and TCER-1 jointly (Joint Downregulated: 36% Overlap, p < 2.094e-13) or DAF-16 alone (Specific Downregulated: 18%, p < 3.791e-08) or TCER-1 alone (Specific Downregulated: 12%, p < 2.213e-04) or downregulated by NHR-49 by oppositely impacted by DAF-16 vs. TCER-1 targets (Oppositely regulated: 17%, p < 0.005).

**Supplementary Figure 2:**
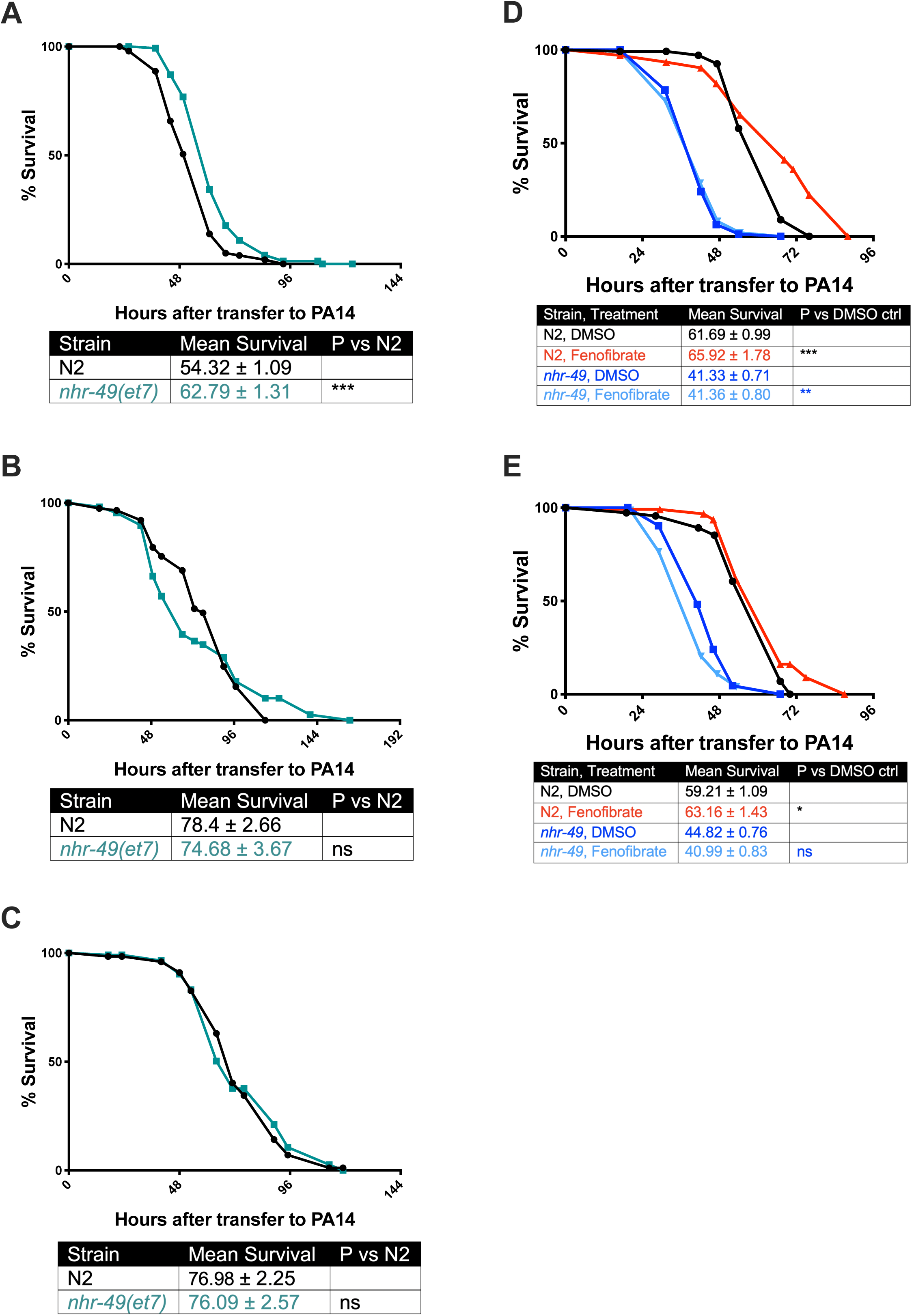
**(A-C)** Survival of L4 staged wild-type animals (N2, black) and nhr-49(et7) mutants (olive) on PA14. (**D, E)** Survival on DMSO- or Fenofibrate-supplemented PA14 plates of L4 staged wild-type animals (N2) and *nhr-49* mutants grown on control DMSO or Fenofibrate from egg-stage onwards. In all panels, data from one independent trial is depicted with mean survival, numbers tested and statistical significance shown in the table associated with each panel.

## List of Supplementary Tables

**Table S1:** Genes differentially expressed between *glp-1* and *nhr-49;glp-1* mutants identified by RNA-Seq

**Table S2:** NHR-49 UP targets essential for *glp-1* mutants’ longevity.

| Wormbase Gene ID | Public Name | Overlapping Group | Avg. % Suppression of Lifespan |
| --- | --- | --- | --- |
| WBGene00000975 | *dhs-12* | DAF-16 UP | -17 |
| WBGene00003473 | *mtl-1* | DAF-16 UP | -22 |
| WBGene00012404 | *Y6G8.2* | DAF-16 UP | -18 |
| WBGene00014030 | *glb-1* | DAF-16 UP | -15 |
| WBGene00021491 | *comt-4* | DAF-16 UP | -11 |
| WBGene00000982 | *dhs-19* | TCER-1 UP | -22 |
| WBGene00000986 | *dhs-23* | TCER-1 UP | -20 |
| WBGene00000988 | *dhs-25* | TCER-1 UP | -21 |
| WBGene00007362 | *cyp-35C1* | TCER-1 UP | -17 |
| WBGene00007848 | *cytb-5.1* | TCER-1 UP | -21 |
| WBGene00007963 | *cyp-25A1* | TCER-1 UP | -10 |
| WBGene00008499 | *cyp-37A1* | TCER-1 UP | -22 |
| WBGene00010659 | *K08D8.5* | TCER-1 UP | -15 |
| WBGene00011830 | *cyp-29A2* | TCER-1 UP | -10 |
| WBGene00013594 | *Y872A.2* | TCER-1 UP | -15 |
| WBGene00015400 | *cyp-35A2* | TCER-1 UP | -16 |
| WBGene00015692 | *ugt-25* | TCER-1 UP | -14 |
| WBGene00016786 | *cyp-35A4* | TCER-1 UP | -20 |
| WBGene00018958 | *cest-27* | TCER-1 UP | -11 |
| WBGene00019473 | *cyp-35A5* | TCER-1 UP | -13 |
| WBGene00000966 | *dhs-2* | DAF-16 & TCER-1 UP | -14 |
| WBGene00000981 | *dhs-18* | DAF-16 & TCER-1 UP | -12 |
| WBGene00001387 | *far-3* | DAF-16 & TCER-1 UP | -12 |
| WBGene00001500 | *ftn-1* | DAF-16 & TCER-1 UP | -18 |
| WBGene00001777 | *gst-29* | DAF-16 & TCER-1 UP | -15 |
| WBGene00004818 | *skr-12* | DAF-16 & TCER-1 UP | -22 |
| WBGene00004994 | *spp-9* | DAF-16 & TCER-1 UP | -12 |
| WBGene00004997 | *spp-12* | DAF-16 & TCER-1 UP | -14 |
| WBGene00007140 | *cyp-29A4* | DAF-16 & TCER-1 UP | -15 |
| WBGene00010706 | *cyp-14A2* | DAF-16 & TCER-1 UP | -17 |
| WBGene00011362 | *cest-1.1* | DAF-16 & TCER-1 UP | -19 |
| WBGene00013904 | *ugt-6* | DAF-16 & TCER-1 UP | -21 |
| WBGene00017390 | *F12A10.1* | DAF-16 & TCER-1 UP | -11 |
| WBGene00020413 | *T10E9.3* | DAF-16 & TCER-1 UP | -22 |
| WBGene00020909 | *W01A11.1* | DAF-16 & TCER-1 UP | -17 |

**Table S3:**
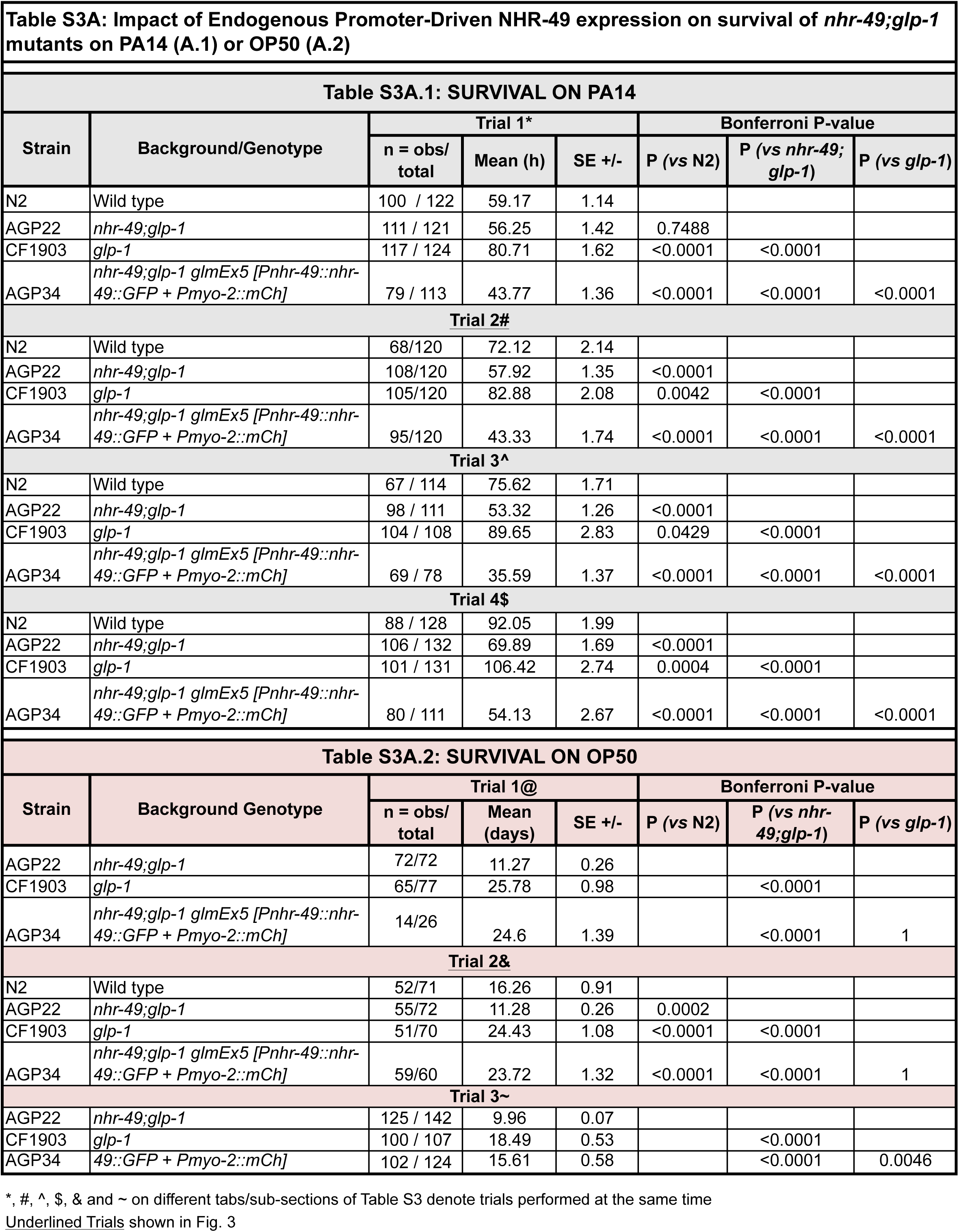
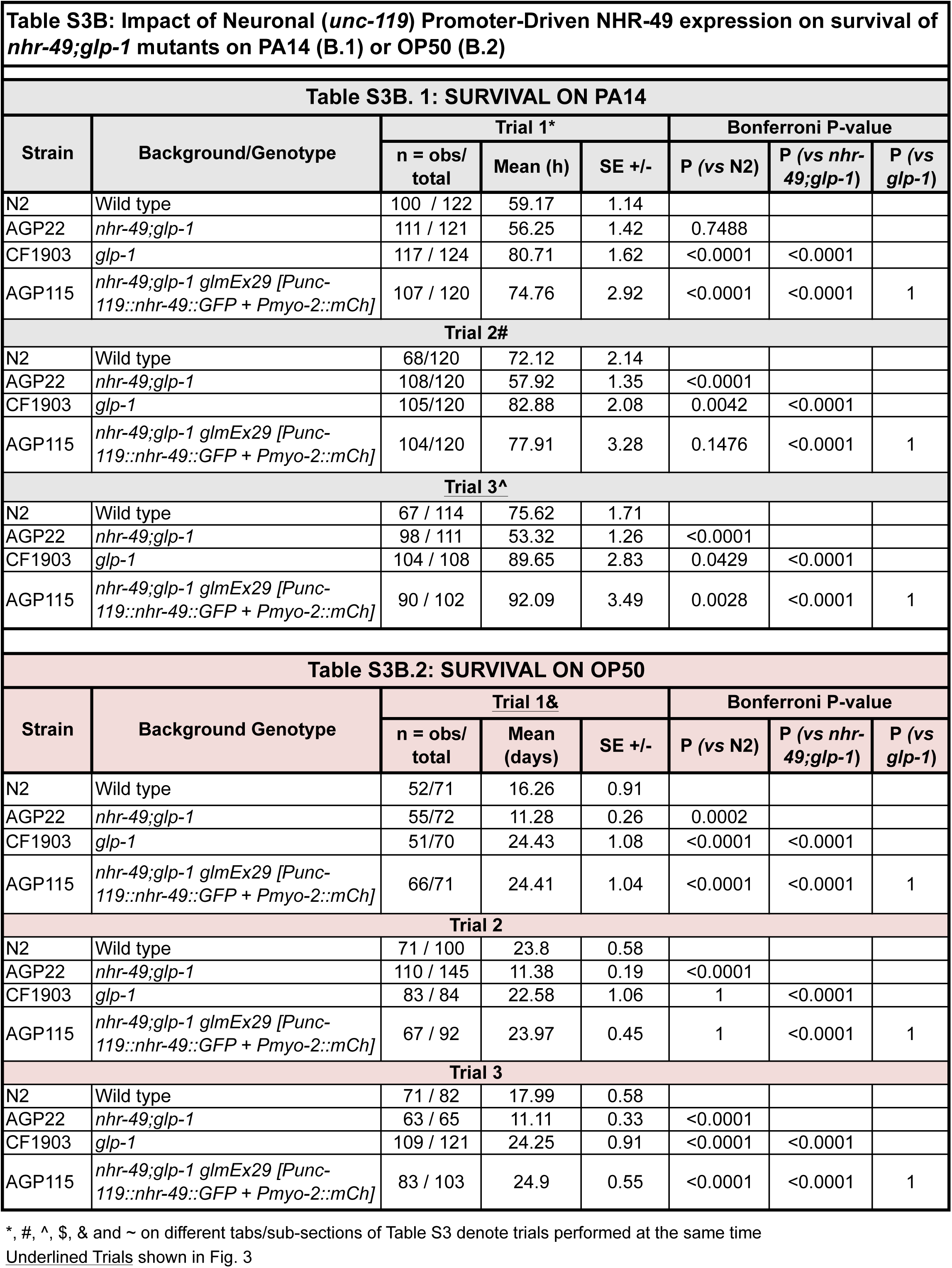
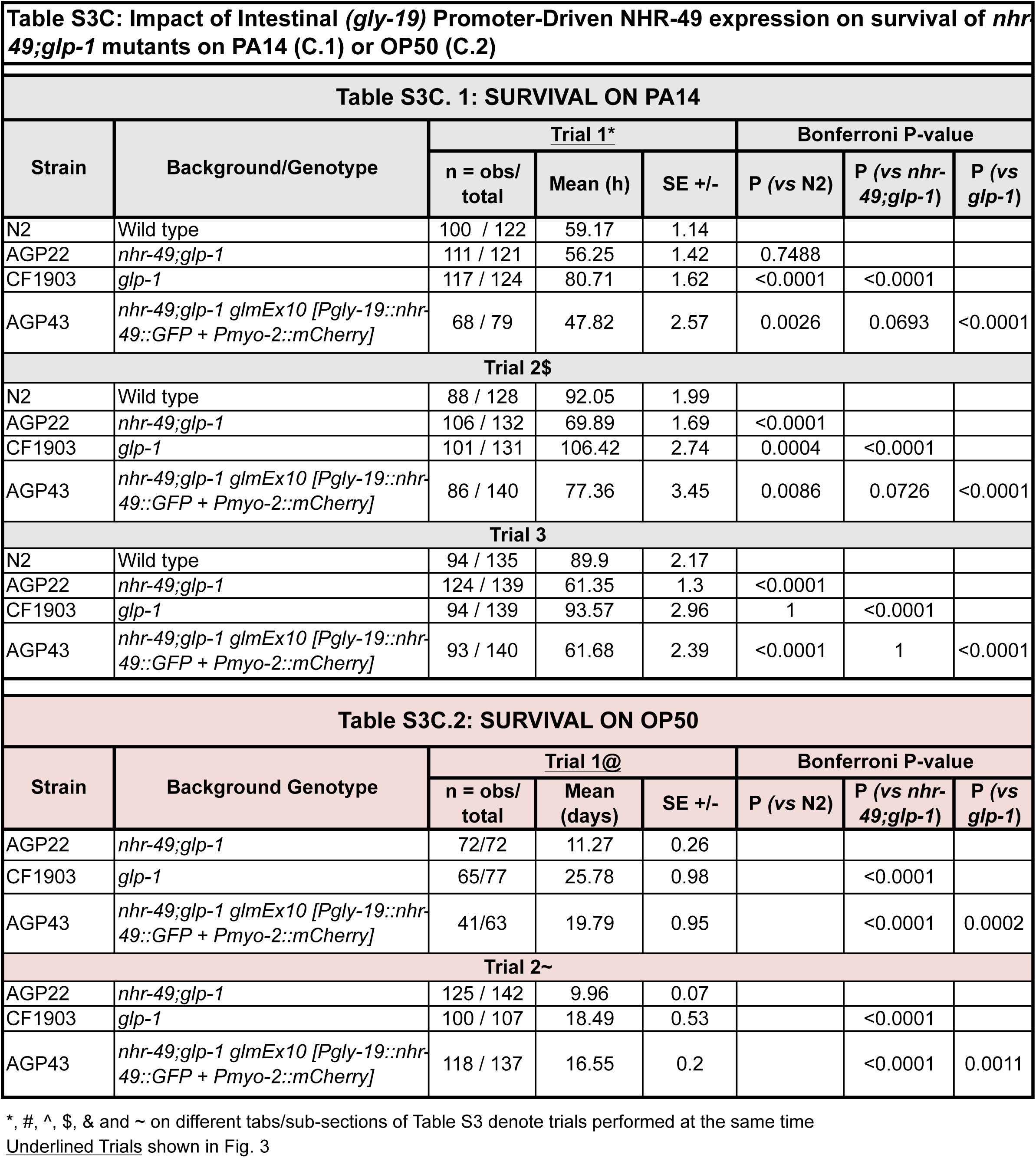
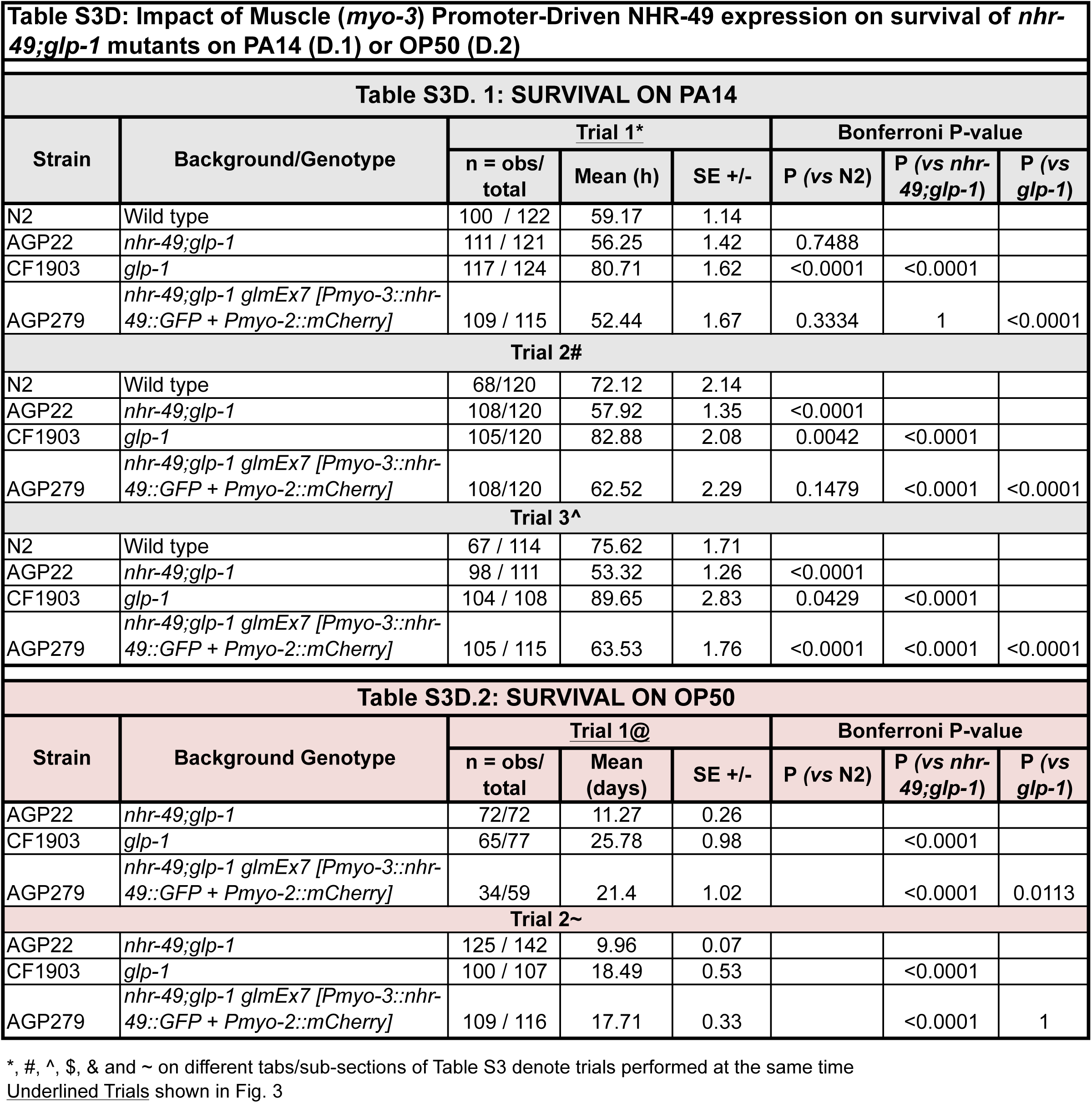
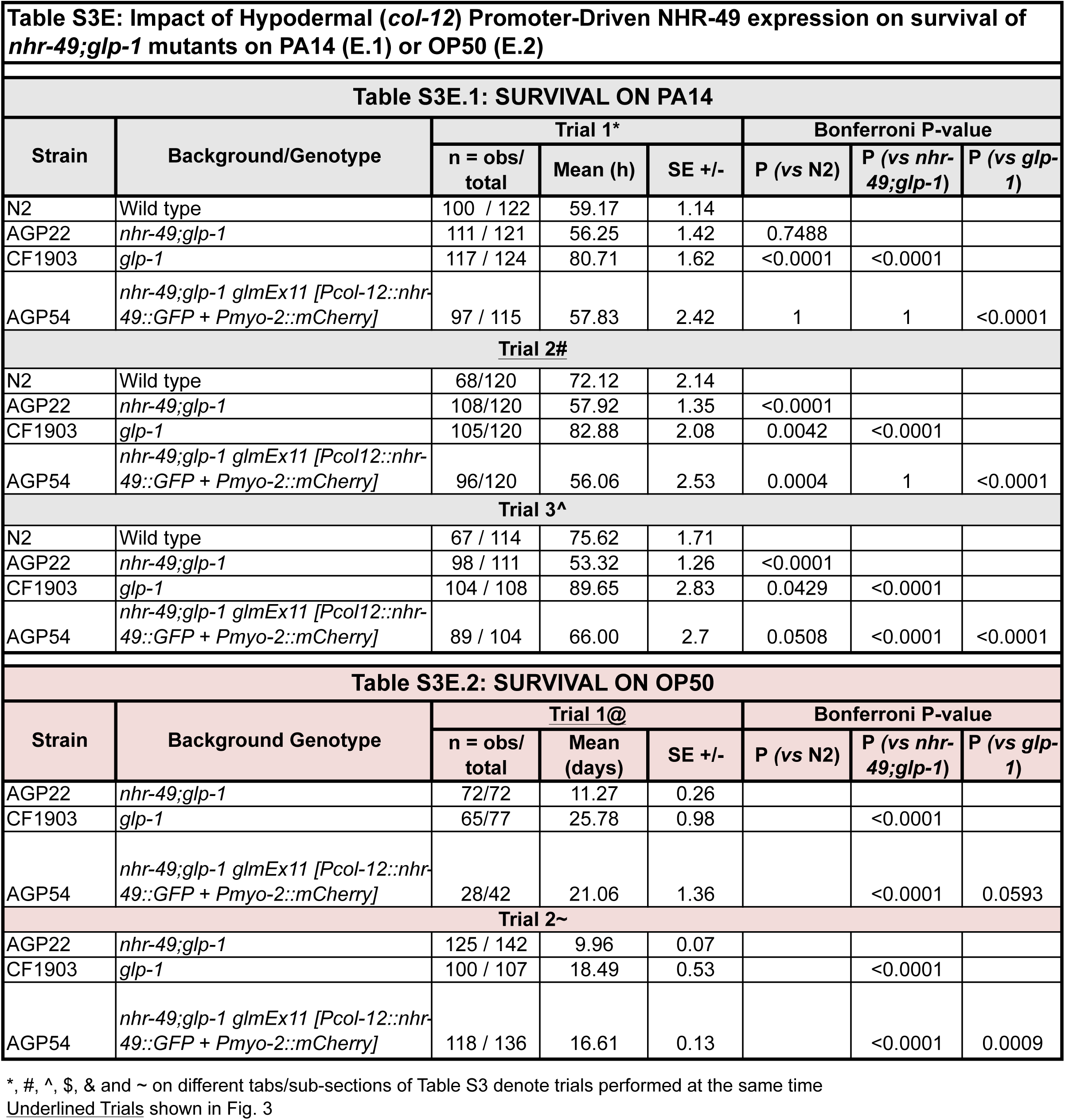
Rescue of *nhr-49;glp-1* mutants PA14 sensitivity and short lifespan on OP50 by NHR-49 expression via endogenous promoter (A) or expression in Neurons (B), Intestine (C), Muscles (D) or Hypodermis (E).

**Table S4:** Rescue of *nhr-49* mutants PA14 sensitivity and short lifespan on OP50 by NHR-49 expression via endogenous promoter (A) or expression in Neurons (B), Intestine (C), Muscles (D) or Hypodermis (E).

| Strain | Background/Genotype | Trial 1* |  |  | Bonferroni P-value |  |
| --- | --- | --- | --- | --- | --- | --- |
|  |  | n = obs/<br>total | Mean (h) | SE +/- | P (vs N2) | P (vs<br>*nhr-49*) |
| N2 | Wild type | 125 / 155 | 67.25 | 1.86 |  |  |
| AGP12a | *nhr-49* | 145 / 167 | 51.89 | 1.41 | <0.0001 |  |
| AGP33 | *nhr-49 glmEx5 [Pnhr-49::*nhr-49::GFP* + *Pmyo-2::mCh*]* | 115 / 131 | 46.68 | 2.12 | <0.0001 | 0.506 |
| **Trial 2#** |  |  |  |  |  |  |
| N2 | Wild type | 93/150 | 68.58 | 2.05 |  |  |
| AGP12a | *nhr-49* | 126/150 | 64.63 | 1.81 | 0.6022 |  |
| AGP33 | *nhr-49 glmEx5 [Pnhr-49::*nhr-49::GFP* + *Pmyo-2::mCh*]* | 87/100 | 74.96 | 3.35 | 0.7647 | 0.0101 |
| **Trial 3^** |  |  |  |  |  |  |
| N2 | Wild type | 63/125 | 78.24 | 2.57 |  |  |
| AGP12a | *nhr-49* | 99/127 | 58 | 1.44 | <0.0001 |  |
| AGP33 | *nhr-49 glmEx5 [Pnhr-49::*nhr-49::GFP* + *Pmyo-2::mCh*]* | 87/127 | 94.98 | 3.08 | 0.002 | <0.0001 |
| **Trial 4\$** | | | | | | |
| N2 | Wild type | 55/90 | 84.82 | 2.63 |  |  |
| AGP12a | *nhr-49* | 102/113 | 60.24 | 1.52 | 0.0001 |  |
| AGP33 | *nhr-49 glmEx5 [Pnhr-49::*nhr-49::GFP* + *Pmyo-2::mCh*]* | 100/125 | 84.35 | 2.17 | 1 | 0.0001 |

| Strain | Background/Genotype | Trial 1@ |  |  | Bonferroni P-value |  |
| --- | --- | --- | --- | --- | --- | --- |
|  |  | n = obs/<br>total | Mean (h) | SE +/- | P (vs N2) | P (vs<br>*nhr-49*) |
| N2 | Wild type | 67/73 | 16.67 | 0.65 |  |  |
| AGP12a | *nhr-49* | 33/77 | 14.39 | 0.69 | 0.1514 |  |
| AGP33 | *nhr-49 glmEx5 [Pnhr-49::*nhr-49::GFP* + *Pmyo-2::mCh*]* | 56/69 | 18.99 | 0.71 | 0.1646 | 0.0001 |
| **Trial 2&** |  |  |  |  |  |  |
| N2 | Wild type | 89 / 122 | 15.42 | 0.49 |  |  |
| AGP12a | *nhr-49* | 110 / 119 | 12.5 | 0.36 | <0.0001 |  |
| AGP33 | *nhr-49 glmEx5 [Pnhr-49::*nhr-49::GFP* + *Pmyo-2::mCh*]* | 97 / 112 | 18 | 0.57 | 0.0424 | <0.0001 |

\*, #, ^, \$, & on different tabs/sub-sections of Table S4 denote trials performed at the same time Underlined Trials shown in Fig.4

| **Table S4B. 1: SURVIVAL ON PA14** |  |  |  |  |  |  |
| --- | --- | --- | --- | --- | --- | --- |
| Strain | Background/Genotype | Trial 1* |  |  | Bonferroni P-value |  |
|  |  | n = obs/<br>total | Mean (h) | SE +/- | P (vs N2) | P (vs<br>*nhr-49*) |
| N2 | Wild type | 125 / 155 | 67.25 | 1.86 |  |  |
| AGP12a | *nhr-49* | 145 / 167 | 51.89 | 1.41 | <0.0001 |  |
| AGP103a | *nhr-49 glmEx20 [Punc-119::*nhr-49::GFP* + *Pmyo-2::mCh*]* | 113 / 124 | 57.87 | 2.82 | 0.9996 | 0.0017 |
| **Trial 2#** |  |  |  |  |  |  |
| N2 | Wild type | 93/150 | 68.58 | 2.05 |  |  |
| AGP12a | *nhr-49* | 126/150 | 64.63 | 1.81 | 0.6022 |  |
| AGP103a | *nhr-49 glmEx20 [Punc-119::*nhr-49::GFP* + *Pmyo-2::mCh*]* | 86/130 | 77.42 | 3.34 | <0.0001 | 0.0002 |
| **Trial 3^** |  |  |  |  |  |  |
| N2 | Wild type | 63/125 | 78.24 | 2.57 |  |  |
| AGP12a | *nhr-49* | 99/127 | 58 | 1.44 | <0.0001 |  |
| AGP103a | *nhr-49 glmEx20 [Punc-119::*nhr-49::GFP* + *Pmyo-2::mCh*]* | 96/125 | 81.54 | 3.19 | 0.7738 | <0.0001 |
| **Trial 4\$** | | | | | | |
| N2 | Wild type | 55/90 | 84.82 | 2.63 |  |  |
| AGP12a | *nhr-49* | 102/113 | 60.24 | 1.52 | 0.0001 |  |
| AGP103a | *nhr-49 glmEx20 [Punc-119::*nhr-49::GFP* + *Pmyo-2::mCh*]* | 95/110 | 84.44 | 2.73 | 1 | 0.0001 |
| **Table S4B.2: SURVIVAL ON OP50** |  |  |  |  |  |  |
| Strain | Background/Genotype | Trial 1@ |  |  | Bonferroni P-value |  |
|  |  | n = obs/<br>total | Mean (h) | SE +/- | P (vs N2) | P (vs<br>*nhr-49*) |
| N2 | Wild type | 67/73 | 16.67 | 0.65 |  |  |
| AGP12a | *nhr-49* | 33/77 | 14.39 | 0.69 | 0.1514 |  |
| AGP103a | *nhr-49 glmEx20 [Punc-119::*nhr-49::GFP* + *Pmyo-2::mCh*]* | 48/57 | 14.58 | 0.63 | 0.1326 | 1 |
| **Trial 2** |  |  |  |  |  |  |
| N2 | Wild type | 74 / 91 | 24.2 | 0.55 |  |  |
| AGP12a | *nhr-49* | 70 / 75 | 14.29 | 0.22 | <0.0001 |  |
| AGP103a | *nhr-49 glmEx20 [Punc-119::*nhr-49::GFP* + *Pmyo-2::mCh*]* | 87 / 95 | 13.98 | 0.2 | <0.0001 | 1 |
| **Trial 3** |  |  |  |  |  |  |
| N2 | Wild type | 71 / 100 | 23.8 | 0.58 |  |  |
| AGP12a | *nhr-49* | 41 / 72 | 9.23 | 0.31 | <0.0001 |  |
| AGP103a | *nhr-49 glmEx20 [Punc-119::*nhr-49::GFP* + *Pmyo-2::mCh*]* | 73 / 91 | 15.37 | 0.57 | <0.0001 | <0.0001 |
| **Trial 4** |  |  |  |  |  |  |
| N2 | Wild type | 71 / 82 | 17.99 | 0.58 |  |  |
| AGP12a | *nhr-49* | 63 / 65 | 11.11 | 0.33 | <0.0001 |  |
| AGP103a | *nhr-49 glmEx20 [Punc-119::*nhr-49::GFP* + *Pmyo-2::mCh*]* | 33 / 76 | 22.23 | 0.71 | 0.0006 | 0.0002 |

\*, #, ^, \$, & on different tabs/sub-sections of Table S4 denote trials performed at the same time

| **Table S4C. 1: SURVIVAL ON PA14** |  |  |  |  |  |  |
| --- | --- | --- | --- | --- | --- | --- |
| Strain | Background/Genotype | Trial 1* |  |  | Bonferroni P-value |  |
|  |  | n = obs/<br>total | Mean (h) | SE +/- | P (vs N2) | P (vs<br>*nhr-49*) |
| N2 | Wild type | 125 / 155 | 67.25 | 1.86 |  |  |
| AGP12a | *nhr-49* | 145 / 167 | 51.89 | 1.41 | <0.0001 |  |
| AGP65 | *nhr-49 glmEx9 [Pgly-19::*nhr-49::GFP* + *Pmyo-2::mCh*]* | 107 / 132 | 50.82 | 2.14 | <0.0001 | 1 |
| **Trial 2#** |  |  |  |  |  |  |
| N2 | Wild type | 93/150 | 68.58 | 2.05 |  |  |
| AGP12a | *nhr-49* | 126/150 | 64.63 | 1.81 | 0.6022 |  |
| AGP65 | *nhr-49 glmEx9 [Pgly-19::*nhr-49::GFP* + *Pmyo-2::mCh*]* | 89/150 | 73.78 | 2.92 | 1 | 0.0201 |
| **Trial 3^** |  |  |  |  |  |  |
| N2 | Wild type | 63/125 | 78.24 | 2.57 |  |  |
| AGP12a | *nhr-49* | 99/127 | 58 | 1.44 | <0.0001 |  |
| AGP65 | *nhr-49 glmEx9 [Pgly-19::*nhr-49::GFP* + *Pmyo-2::mCh*]* | 87/125 | 77.6 | 2.73 | 1 | <0.0001 |
| **Trial 4\$** | | | | | | |
| N2 | Wild type | 55/90 | 84.82 | 2.63 |  |  |
| AGP12a | *nhr-49* | 102/113 | 60.24 | 1.52 | 0.0001 |  |
| AGP65 | *nhr-49 glmEx9 [Pgly-19::*nhr-49::GFP* + *Pmyo-2::mCh*]* | 100/116 | 75.5 | 2.37 | 0.2823 | 0.0001 |
| **Table S4C.2: SURVIVAL ON OP50** |  |  |  |  |  |  |
| Strain | Background/Genotype | Trial 1@ |  |  | Bonferroni P-value |  |
|  |  | n = obs/<br>total | Mean (h) | SE +/- | P (vs N2) | P (vs<br>*nhr-49*) |
| N2 | Wild type | 67/73 | 16.67 | 0.65 |  |  |
| AGP12a | *nhr-49* | 33/77 | 14.39 | 0.69 | 0.1514 |  |
| AGP65 | *nhr-49 glmEx9 [Pgly-19::*nhr-49::GFP* + *Pmyo-2::mCh*]* | 47/56 | 15.85 | 0.71 | 1 | 1 |
| **Trial 2&** |  |  |  |  |  |  |
| N2 | Wild type | 89 / 122 | 15.42 | 0.49 |  |  |
| AGP12a | *nhr-49* | 110 / 119 | 12.5 | 0.36 | <0.0001 |  |
| AGP65 | *nhr-49 glmEx9 [Pgly-19::*nhr-49::GFP* + *Pmyo-2::mCh*]* | 97 / 118 | 15.25 | 0.4 | 1 | <0.0001 |

\*, #, ^, \$, & on different tabs/sub-sections of Table S4 denote trials performed at the same time Underlined Trials shown in Fig.4

| **Table S4D. 1: SURVIVAL ON PA14** |  |  |  |  |  |  |
| --- | --- | --- | --- | --- | --- | --- |
| Strain | Background/Genotype | Trial 1* |  |  | Bonferroni P-value |  |
|  |  | n = obs/<br>total | Mean (h) | SE +/- | P (vs N2) | P (vs *nhr-49*) |
| N2 | Wildtype | 125 / 155 | 67.25 | 1.86 |  |  |
| AGP12a | *nhr-49* | 145 / 167 | 51.89 | 1.41 | <0.0001 |  |
| AGP63 | *nhr-49 glmEx8 [Pmyo-3::*nhr-49::GFP* + *Pmyo-2::mCh*]* | 137 / 153 | 44.92 | 1.55 | <0.0001 | 0.0819 |
| **Trial 2#** |  |  |  |  |  |  |
| N2 | Wildtype | 93/150 | 68.58 | 2.05 |  |  |
| AGP12a | *nhr-49* | 126/150 | 64.63 | 1.81 | 0.6022 |  |
| AGP63 | *nhr-49 glmEx8 [Pmyo-3::*nhr-49::GFP* + *Pmyo-2::mCh*]* | 115/150 | 65.11 | 1.73 | 1 | 1 |
| **Trial 3^** |  |  |  |  |  |  |
| N2 | Wildtype | 63/125 | 78.24 | 2.57 |  |  |
| AGP12a | *nhr-49* | 99/127 | 58 | 1.44 | <0.0001 |  |
| AGP63 | *nhr-49 glmEx8 [Pmyo-3::*nhr-49::GFP* + *Pmyo-2::mCh*]* | 91/126 | 68.06 | 1.59 | 0.0124 | <0.0001 |
| **Trial 4\$** | | | | | | |
| N2 | Wildtype | 55/90 | 84.82 | 2.63 |  |  |
| AGP12a | *nhr-49* | 102/113 | 60.24 | 1.52 | 0.0001 |  |
| AGP63 | *nhr-49 glmEx8 [Pmyo-3::*nhr-49::GFP* + *Pmyo-2::mCh*]* | 105/119 | 65.73 | 1.27 | 0.0001 | 0.0991 |
| **Table S4D.2: SURVIVAL ON OP50** |  |  |  |  |  |  |
| Strain | Background/Genotype | Trial 1@ |  |  | Bonferroni P-value |  |
|  |  | n = obs/<br>total | Mean (h) | SE +/- | P (vs N2) | P (vs *nhr-49*) |
| N2 | Wildtype | 67/73 | 16.67 | 0.65 |  |  |
| AGP12a | *nhr-49* | 33/77 | 14.39 | 0.69 | 0.1514 |  |
| AGP63 | *nhr-49 glmEx8 [Pmyo-3::*nhr-49::GFP* + *Pmyo-2::mCh*]* | 73/75 | 11.84 | 0.54 | 0.0001 | 0.0893 |
| **Trial 2&** |  |  |  |  |  |  |
| N2 | Wildtype | 89 / 122 | 15.42 | 0.49 |  |  |
| AGP12a | *nhr-49* | 110 / 119 | 12.5 | 0.36 | <0.0001 |  |
| AGP63 | *nhr-49 glmEx8 [Pmyo-3::*nhr-49::GFP* + *Pmyo-2::mCh*]* | 75 / 80 | 12.01 | 0.71 | 0.0001 | 1 |

\*, #, ^, \$, & on different tabs/sub-sections of Table S4 denote trials performed at the same time Underlined Trials shown in Fig.4

| Strain | Background/Genotype | Trial 1* |  |  | Bonferroni P-value |  |
| --- | --- | --- | --- | --- | --- | --- |
|  |  | n = obs/<br>total | Mean (h) | SE +/- | P (vs N2) | P (vs<br>*nhr-49*) |
| N2 | Wild type | 125 / 155 | 67.25 | 1.86 |  |  |
| AGP12a | *nhr-49* | 145 / 167 | 51.89 | 1.41 | <0.0001 |  |
| AGP53 | *nhr-49 glmEx11 [Pcol-12::*nhr-49::GFP* + *Pmyo-2::mCh*]* | 138 / 150 | 30.21 | 1.77 | <0.0001 | <0.0001 |
| **Trial 2#** |  |  |  |  |  |  |
| N2 | Wild type | 93/150 | 68.58 | 2.05 |  |  |
| AGP12a | *nhr-49* | 126/150 | 64.63 | 1.81 | 0.6022 |  |
| AGP63 | *nhr-49 glmEx8 [Pmyo-3::*nhr-49::GFP* + *Pmyo-2::mCh*]* | 90/128 | 48.55 | 2.56 | <0.0001 | 0.0001 |
| **Trial 3^** |  |  |  |  |  |  |
| N2 | Wild type | 63/125 | 78.24 | 2.57 |  |  |
| AGP12a | *nhr-49* | 99/127 | 58 | 1.44 | <0.0001 |  |
| AGP53 | *nhr-49 glmEx11 [Pcol-12::*nhr-49::GFP* + *Pmyo-2::mCh*]* | 79/100 | 46.27 | 2.72 | <0.0001 | 0.0415 |
| **Trial 4\$** | | | | | | |
| N2 | Wild type | 55/90 | 84.82 | 2.63 |  |  |
| AGP12a | *nhr-49* | 102/113 | 60.24 | 1.52 | 0.0001 |  |
| AGP53 | *nhr-49 glmEx11 [Pcol-12::*nhr-49::GFP* + *Pmyo-2::mCh*]* | 28/32 | 48.62 | 4.65 | 0.0001 | 0.146 |

| Strain | Background/Genotype | Trial 1@ |  |  | Bonferroni P-value |  |
| --- | --- | --- | --- | --- | --- | --- |
|  |  | n = obs/<br>total | Mean (h) | SE +/- | P (vs N2) | P (vs<br>*nhr-49*) |
| N2 | Wild type | 67/73 | 16.67 | 0.65 |  |  |
| AGP12a | *nhr-49* | 33/77 | 14.39 | 0.69 | 0.1514 |  |
| AGP53 | *nhr-49 glmEx11 [Pcol-12::*nhr-49::GFP* + *Pmyo-2::mCh*]* | 59/70 | 16.89 | 0.65 | 1 | 0.0926 |
| **Trial 2&** |  |  |  |  |  |  |
| N2 | Wild type | 89 / 122 | 15.42 | 0.49 |  |  |
| AGP12a | *nhr-49* | 110 / 119 | 12.5 | 0.36 | <0.0001 |  |
| AGP53 | *nhr-49 glmEx11 [Pcol-12::*nhr-49::GFP* + *Pmyo-2::mCh*]* | 131 / 141 | 14.86 | 0.45 | 1 | <0.0001 |

\*, #, ^, \$, & on different tabs/sub-sections of Table S4 denote trials performed at the same time Underlined Trials shown in Fig.4

**Table S5:** Impact on wild-type worms’ survival on PA14 and lifespan on OP50 by NHR-49 over expression via endogenous promoter (A) or expression in Neurons (B), Intestine (C), Muscles (D) or Hypodermis (E).

| Table S5A. 1: SURVIVAL ON PA14 |  |  |  |  |  |  |
| --- | --- | --- | --- | --- | --- | --- |
| Strain | Background/Genotype | Trial 1* |  |  | Bonferroni P-value | Non-Bonferroni P-value |
|  |  | n = obs/total | Mean (h) | SE +/- |  |  |
| N2 | Wild type | 126 / 145 | 54.32 | 1.09 |  |  |
| AGP24f | *glmEx5 [Pnhr-49::nhr-49::GFP + Pmyo-2::mCh]* | 123 / 153 | 68.14 | 1.7 | <0.0001 | <0.0001 |
| Trial 2# |  |  |  |  |  |  |
| N2 | Wild type | 60 / 117 | 78.4 | 2.66 |  |  |
| AGP24f | *glmEx5 [Pnhr-49::nhr-49::GFP + Pmyo-2::mCh]* | 61 / 110 | 71.05 | 3.78 | 0.229 | 0.0286 |
| Trial 3^ |  |  |  |  |  |  |
| N2 | Wild type | 102 / 129 | 76.98 | 2.25 |  |  |
| AGP24f | *glmEx5 [Pnhr-49::nhr-49::GFP + Pmyo-2::mCh]* | 75 / 121 | 76.89 | 3.06 | 1 | 0.895 |
| Table S5A.2: SURVIVAL ON OP50 |  |  |  |  |  |  |
| Strain | Background/Genotype | Trial 1@ |  |  | Bonferroni P-value (vs N2) | Non-Bonferroni P (vs N2) |
|  |  | n = obs/total | Mean (h) | SE +/- |  |  |
| N2 | Wild type | 67 / 73 | 16.67 | 0.65 |  |  |
| AGP12a | *nhr-49* | 33 / 77 | 14.39 | 0.69 | 0.1514 |  |
| AGP24f | *glmEx5 [Pnhr-49::nhr-49::GFP + Pmyo-2::mCh]* | 35 / 44 | 20.83 | 1.01 | 0.0109 | 0.0016 |
| Trial 2~ |  |  |  |  |  |  |
| N2 | Wild type | 73 / 121 | 17.81 | 0.56 |  |  |
| AGP12a | *nhr-49* | 69 / 76 | 14.6 | 0.54 | 0.0003 |  |
| AGP24f | *glmEx5 [Pnhr-49::nhr-49::GFP + Pmyo-2::mCh]* | 70 / 115 | 17.58 | 0.75 | 1 | 0.603 |

\*, #, ^, &, @, ~ on different tabs/sub-sections of Table S5 denote trials performed at the same time Underlined Trials shown in Fig. 5

| Strain | Background/Genotype | Trial 1* |  |  | Bonferroni P-value | Non-Bonferroni P-value |
| --- | --- | --- | --- | --- | --- | --- |
|  |  | n = obs/total | Mean (h) | SE +/- |  |  |
| N2 | Wild type | 126 / 145 | 54.32 | 1.09 |  |  |
| AGP102 | *glmEx20 [Pnhr-49::unc-119::GFP + Pmyo-2::mCh]* | 114 / 136 | 73.72 | 1.61 | <0.0001 | <0.0001 |
| **Trial 2#** |  |  |  |  |  |  |
| N2 | Wild type | 60 / 117 | 78.4 | 2.66 |  |  |
| AGP102 | *glmEx20 [Pnhr-49::unc-119::GFP + Pmyo-2::mCh]* | 88/120 | 70.79 | 2.11 | 0.093 | 0.0116 |
| **Trial 3^** |  |  |  |  |  |  |
| N2 | Wild type | 102 / 129 | 76.98 | 2.25 |  |  |
| AGP102 | *glmEx20 [Pnhr-49::unc-119::GFP + Pmyo-2::mCh]* | 96/127 | 86.37 | 2.44 | 0.0333 | 0.0135 |

| Strain | Background/Genotype | Trial 1@ |  |  | Bonferroni P-value | Non-Bonferroni P-value |
| --- | --- | --- | --- | --- | --- | --- |
|  |  | n = obs/total | Mean (h) | SE +/- |  |  |
| N2 | Wild type | 67 / 73 | 16.67 | 0.65 |  |  |
| AGP102 | *glmEx20 [Pnhr-49::unc-119::GFP + Pmyo-2::mCh]* | 39/57 | 19.31 | 0.75 | 0.0651 | 0.0093 |
| **Trial 2** |  |  |  |  |  |  |
| N2 | Wild type | 74 / 91 | 24.2 | 0.55 |  |  |
| AGP102 | *glmEx20 [Pnhr-49::unc-119::GFP + Pmyo-2::mCh]* | 57 / 94 | 22.35 | 0.6 | 0.4163 | 0.0416 |
| **Trial 3** |  |  |  |  |  |  |
| N2 | Wild type | 71 / 100 | 23.8 | 0.58 |  |  |
| AGP102 | *glmEx20 [Pnhr-49::unc-119::GFP + Pmyo-2::mCh]* | 40 / 71 | 21.53 | 0.81 | 0.074 | 0.0092 |
| **Trial 4** |  |  |  |  |  |  |
| N2 | Wildtype | 71 / 82 | 17.99 | 0.58 |  |  |
| AGP102 | *glmEx20 [Pnhr-49::unc-119::GFP + Pmyo-2::mCh]* | 32 / 70 | 23.23 | 0.97 | <0.0001 | <0.0001 |

\*, #, ^, &, @, ~ on different tabs/sub-sections of Table S5 denote trials performed at the same time Underlined Trials shown in Fig. 5

| Table S5C. 1: SURVIVAL ON PA14 |  |  |  |  |  |  |
| --- | --- | --- | --- | --- | --- | --- |
| Strain | Background/Genotype | Trial 1* |  |  | Bonferroni P-value | Non-Bonferroni P-value |
|  |  | n = obs/ total | Mean (h) | SE +/- |  |  |
| N2 | Wild type | 126 / 145 | 54.32 | 1.09 |  |  |
| AGP40 | *glmEx9 [Pgly-19::nhr-49::GFP + Pmyo-2::mCherry]* | 117 / 131 | 73.6 | 1.83 | <0.0001 | <0.0001 |
| Trial 2# |  |  |  |  |  |  |
| N2 | Wild type | 60/117 | 78.4 | 2.66 |  |  |
| AGP40 | *glmEx9 [Pgly-19::nhr-49::GFP + Pmyo-2::mCherry]* | 92/128 | 94.83 | 2.72 | 0.0007 | 0.0001 |
| Trial 3^ |  |  |  |  |  |  |
| N2 | Wildtype | 102/129 | 76.98 | 2.25 |  |  |
| AGP40 | *glmEx9 [Pgly-19::nhr-49::GFP + Pmyo-2::mCherry]* | 96/124 | 91.97 | 2.65 | 0.0002 | 0.0001 |
| Table S5C.2: SURVIVAL ON OP50 |  |  |  |  |  |  |
| Strain | Background Genotype | Trial 1@ |  |  | Bonferroni P-value (vs N2) | Non-Bonferroni P (vs N2) |
|  |  | n = obs/ total | Mean (days) | SE +/- |  |  |
| N2 | Wild type | 67/73 | 16.67 | 0.65 |  |  |
| AGP12a | *nhr-49* | 33/77 | 14.39 | 0.69 | 0.1514 |  |
| AGP40 | *glmEx9 [Pgly-19::nhr-49::GFP + Pmyo-2::mCherry]* | 64/67 | 18.49 | 0.82 | 0.6491 | 0.0927 |
| Trial 2~ |  |  |  |  |  |  |
| N2 | Wild type | 73 / 121 | 17.81 | 0.56 |  |  |
| AGP12a | *nhr-49* | 69 / 76 | 14.6 | 0.54 | 0.0003 |  |
| AGP40 | *glmEx9 [Pgly-19::nhr-49::GFP + Pmyo-2::mCherry]* | 86 / 116 | 19.06 | 0.4 | 1 | 0.202 |

\*, #, ^, &, @, ~ on different tabs/sub-sections of Table S5 denote trials performed at the same time Underlined Trials shown in Fig. 5

| Table S5D. 1: SURVIVAL ON PA14 |  |  |  |  |  |  |
| --- | --- | --- | --- | --- | --- | --- |
| Strain | Background/Genotype | Trial 1* |  |  | Bonferroni P-value | Non-Bonferroni P-value |
|  |  | n = obs/total | Mean (h) | SE +/- |  |  |
| N2 | Wild type | 126 / 145 | 54.32 | 1.09 |  |  |
| AGP27 | *glmEx9 [Pmyo-3::nhr-49::GFP + Pmyo-2::mCherry]* | 106 / 126 | 61.12 | 2.11 | 0.0007 | 0.0001 |
| Trial 2# |  |  |  |  |  |  |
| N2 | Wild type | 60/117 | 78.4 | 2.66 |  |  |
| AGP27 | *glmEx9 [Pmyo-3::nhr-49::GFP + Pmyo-2::mCherry]* | 61/100 | 72.85 | 4.25 | 1 | 0.289 |
| Trial 3^ |  |  |  |  |  |  |
| N2 | Wild type | 102/129 | 76.98 | 2.25 |  |  |
| AGP27 | *glmEx9 [Pmyo-3::nhr-49::GFP + Pmyo-2::mCherry]* | 66/100 | 72.11 | 4.28 | 1 | 0.363 |
| Table S5D.2: SURVIVAL ON OP50 |  |  |  |  |  |  |
| Strain | Background Genotype | Trial 1@ |  |  | Bonferroni P-value (vs N2) | Non-Bonferroni P (vs N2) |
|  |  | n = obs/total | Mean (days) | SE +/- |  |  |
| N2 | Wild type | 67/73 | 16.67 | 0.65 |  |  |
| AGP12a | *nhr-49* | 33/77 | 14.39 | 0.69 | 0.1514 |  |
| AGP27 | *glmEx9 [Pmyo-3::nhr-49::GFP + Pmyo-2::mCherry]* | 39/52 | 14.6 | 0.55 | 0.5331 | 0.0762 |
| Trial 2~ |  |  |  |  |  |  |
| N2 | Wild type | 73 / 121 | 17.81 | 0.56 |  |  |
| AGP12a | *nhr-49* | 69 / 76 | 14.6 | 0.54 | 0.0003 |  |
| AGP27 | *glmEx9 [Pmyo-3::nhr-49::GFP + Pmyo-2::mCherry]* | 84 / 109 | 19.89 | 0.68 | 0.0389 | 0.0065 |

\*, #, ^, &, @, ~ on different tabs/sub-sections of Table S5 denote trials performed at the same time Underlined Trials shown in Fig. 5

| Strain | Background/Genotype | Trial 1* |  |  | Bonferroni P-value | Non-Bonferroni P-value |
| --- | --- | --- | --- | --- | --- | --- |
|  |  | n = obs/total | Mean (h) | SE +/- |  |  |
| N2 | Wild type | 126 / 145 | 54.32 | 1.09 |  |  |
| AGP45 | *glmEx11 [Pcol-12::nhr-49::GFP + Pmyo-2::mCherry]* | 117 / 137 | 77.16 | 1.44 | <0.0001 | <0.0001 |
| **Trial 2#** |  |  |  |  |  |  |
| N2 | Wild type | 60/117 | 78.4 | 2.66 |  |  |
| AGP45 | *glmEx11 [Pcol-12::nhr-49::GFP + Pmyo-2::mCherry]* | 77/125 | 85.91 | 3.67 | 1 | 0.326 |
| **Trial 3^** |  |  |  |  |  |  |
| N2 | Wildtype | 102/129 | 76.98 | 2.25 |  |  |
| AGP45 | *glmEx11 [Pcol-12::nhr-49::GFP + Pmyo-2::mCherry]* | 63/97 | 82.68 | 2.65 | 0.3504 | 0.013 |

| Strain | Background Genotype | Trial 1@ |  |  | Bonferroni P-value (vs N2) | Non-Bonferroni P (vs N2) |
| --- | --- | --- | --- | --- | --- | --- |
|  |  | n = obs/total | Mean (days) | SE +/- |  |  |
| N2 | Wild type | 67/73 | 16.67 | 0.65 |  |  |
| AGP12a | *nhr-49* | 33/77 | 14.39 | 0.69 | 0.1514 |  |
| AGP45 | *glmEx11 [Pcol-12::nhr-49::GFP + Pmyo-2::mCherry]* | 57/66 | 17.27 | 0.67 | 1 | 0.4482 |
| **Trial 2~** |  |  |  |  |  |  |
| N2 | Wild type | 73 / 121 | 17.81 | 0.56 |  |  |
| AGP12a | *nhr-49* | 69 / 76 | 14.6 | 0.54 | 0.0003 |  |
| AGP45 | *glmEx11 [Pcol-12::nhr-49::GFP + Pmyo-2::mCherry]* | 95 / 120 | 18.3 | 0.53 | 1 | 0.468 |

\*, #, ^, &, @, ~ on different tabs/sub-sections of Table S5 denote trials performed at the same time Underlined Trials shown in Fig. 5

**Table S6:** Impact on *glp-1* mutants PA14 resistance and longevity on OP50 by NHR-49 expression via endogenous promoter (A) or expression in Neurons (B), Intestine (C), Muscles (D) or Hypodermis (E).

| Strain | Background/Genotype | Trial 1* |  |  | Bonferroni P-value |  |
| --- | --- | --- | --- | --- | --- | --- |
|  |  | n = obs/<br>total | Mean (h) | SE +/- | P (vs N2) | P (vs<br>*glp-1*) |
| N2 | Wild type | 83 / 105 | 67.31 | 1.37 |  |  |
| CF1903 | *glp-1* | 116 / 127 | 83.68 | 2.04 | <0.0001 |  |
| AGP29 | *glp-1 glmEx6 [Pnhr-49::nhr-49::GFP + Pmyo-2::mCh]* | 105 / 112 | 38.6 | 1.57 | <0.0001 | <0.0001 |
| **Trial 2#** |  |  |  |  |  |  |
| N2 | Wild type | 61/112 | 72.21 | 3.63 |  |  |
| CF1903 | *glp-1* | 95/109 | 88.89 | 2.57 | 0.001 |  |
| AGP29 | *glp-1 glmEx6 [Pnhr-49::nhr-49::GFP + Pmyo-2::mCh]* | 98/115 | 44.15 | 1.92 | <0.0001 | <0.0001 |
| **Trial 3^** |  |  |  |  |  |  |
| N2 | Wild type | 68 / 120 | 68.39 | 2.2 |  |  |
| CF1903 | *glp-1* | 102 / 117 | 94.91 | 3.2 | 0.0001 |  |
| AGP29 | *glp-1 glmEx6 [Pnhr-49::nhr-49::GFP + Pmyo-2::mCh]* | 94 / 105 | 39.35 | 1.56 | 0.0001 | 0.0001 |
| **Trial 4** |  |  |  |  |  |  |
| N2 | Wild type | 88 / 128 | 92.05 | 1.99 |  |  |
| CF1903 | *glp-1* | 101 / 131 | 106.42 | 2.74 | 0.0004 |  |
| AGP29 | *glp-1 glmEx6 [Pnhr-49::nhr-49::GFP + Pmyo-2::mCh]* | 132 / 156 | 53.58 | 2.18 | <0.0001 | <0.0001 |

| Strain | Background/Genotype | Trial 1@ |  |  | Bonferroni P-value |  |
| --- | --- | --- | --- | --- | --- | --- |
|  |  | n = obs/<br>total | Mean (h) | SE +/- | P (vs N2) | P (vs<br>*glp-1*) |
| AGP22 | *nhr-49;glp-1* | 72/72 | 11.27 | 0.26 |  | <0.0001 |
| CF1903 | *glp-1* | 65/77 | 25.78 | 0.98 |  |  |
| AGP29 | *glp-1 glmEx6 [Pnhr-49::nhr-49::GFP + Pmyo-2::mCh]* | 13/14 | 22.62 | 1.85 |  | 0.0755 |
| **Trial 2~** |  |  |  |  |  |  |
| N2 | Wild type | 82 / 122 | 15.53 | 0.58 |  |  |
| CF1903 | *glp-1* | 112 / 124 | 22.98 | 0.67 | <0.0001 |  |
| AGP29 | *glp-1 glmEx6 [Pnhr-49::nhr-49::GFP + Pmyo-2::mCh]* | 94 / 108 | 21.31 | 0.42 | <0.0001 | 0.003 |
| **Trial 3\$** | | | | | | |
| N2 | Wild type | 73 / 121 | 17.81 | 0.56 |  |  |
| CF1903 | *glp-1* | 101 / 114 | 23.35 | 0.69 | <0.0001 |  |
| AGP29 | *glp-1 glmEx6 [Pnhr-49::nhr-49::GFP + Pmyo-2::mCh]* | 55 / 120 | 21.71 | 0.53 | <0.0001 | 0.3082 |

\*, #, ^, &, @, ~ on different tabs/sub-sections of Table S6 denote trials performed at the same time Underlined Trials shown in Fig. 6

| Strain | Background/Genotype | Trial 1* |  |  | Bonferroni P-value |  |
| --- | --- | --- | --- | --- | --- | --- |
|  |  | n = obs/<br>total | Mean (h) | SE +/- | P (vs N2) | P (vs<br>*glp-1*) |
| N2 | Wild type | 83 / 105 | 67.31 | 1.37 |  |  |
| CF1903 | *glp-1* | 116 / 127 | 83.68 | 2.04 | <0.0001 |  |
| AGP105 | *glp-1 glmEx20 [Punc-119::nhr-49::GFP + Pmyo-2::mCh]* | 97 / 117 | 70.81 | 2.96 | 0.0174 | 0.242 |
| **Trial 2#** |  |  |  |  |  |  |
| N2 | Wild type | 61/112 | 72.21 | 3.63 |  |  |
| CF1903 | *glp-1* | 95/109 | 88.89 | 2.57 | 0.001 |  |
| AGP105 | *glp-1 glmEx20 [Punc-119::nhr-49::GFP + Pmyo-2::mCh]* | 111/120 | 59.86 | 3.3 | 0.1603 | <0.0001 |
| **Trial 3^** |  |  |  |  |  |  |
| N2 | Wild type | 68 / 120 | 68.39 | 2.2 |  |  |
| CF1903 | *glp-1* | 102 / 117 | 94.91 | 3.2 | 0.0001 |  |
| AGP105 | *glp-1 glmEx20 [Punc-119::nhr-49::GFP + Pmyo-2::mCh]* | 73 / 80 | 73.22 | 4.93 | 1 | 0.0582 |

| Strain | Background/Genotype | Trial 1 |  |  | Bonferroni P-value |  |
| --- | --- | --- | --- | --- | --- | --- |
|  |  | n = obs/<br>total | Mean (h) | SE +/- | P (vs N2) | P (vs<br>*glp-1*) |
| N2 | Wild type | 52/71 | 16.26 | 0.91 |  |  |
| CF1903 | *glp-1* | 51/70 | 24.43 | 1.08 | <0.0001 |  |
| AGP105 | *glp-1 glmEx20 [Punc-119::nhr-49::GFP + Pmyo-2::mCh]* | 29 / 42 | 21.78 | 1.86 | 0.01 | 1 |
| **Trial 2** |  |  |  |  |  |  |
| N2 | Wild type | 74 / 91 | 24.2 | 0.55 |  |  |
| CF1903 | *glp-1* | 120 / 124 | 30.54 | 0.79 | <0.0001 |  |
| AGP105 | *glp-1 glmEx20 [Punc-119::nhr-49::GFP + Pmyo-2::mCh]* | 70 / 80 | 31.4 | 1.1 | <0.0001 | 1 |
| **Trial 3** |  |  |  |  |  |  |
| N2 | Wild type | 71 / 100 | 23.8 | 0.58 |  |  |
| CF1903 | *glp-1* | 83 / 84 | 22.58 | 1.06 | 1 |  |
| AGP105 | *glp-1 glmEx20 [Punc-119::nhr-49::GFP + Pmyo-2::mCh]* | 51 / 63 | 26.69 | 0.74 | 0.1157 | 1 |
| **Trial 4** |  |  |  |  |  |  |
| N2 | Wild type | 71 / 82 | 17.99 | 0.58 |  |  |
| CF1903 | *glp-1* | 109 / 121 | 24.25 | 0.91 | <0.0001 |  |
| AGP105 | *glp-1 glmEx20 [Punc-119::nhr-49::GFP + Pmyo-2::mCh]* | 99 / 120 | 28.14 | 0.89 | <0.0001 | 0.38 |

\*, #, ^, &, @, ~ on different tabs/sub-sections of Table S6 denote trials performed at the same time Underlined Trials shown in Fig. 6

| Strain | Background/Genotype | Trial 1* |  |  | Bonferroni P-value |  |
| --- | --- | --- | --- | --- | --- | --- |
|  |  | n = obs/<br>total | Mean (h) | SE +/- | P (vs N2) | P (vs *glp-1*) |
| N2 | Wild type | 83 / 105 | 67.31 | 1.37 |  |  |
| CF1903 | *glp-1* | 116 / 127 | 83.68 | 2.04 | <0.0001 |  |
| AGP44 | *glp-1 glmEx10 [Pgly-19::nhr-49::GFP + Pmyo-2::mCh]* | 89 / 108 | 80.91 | 2.67 | <0.0001 | 1 |
| **Trial 2#** |  |  |  |  |  |  |
| N2 | Wild type | 61/112 | 72.21 | 3.63 |  |  |
| CF1903 | *glp-1* | 95/109 | 88.89 | 2.57 | 0.001 |  |
| AGP44 | *glp-1 glmEx10 [Pgly-19::nhr-49::GFP + Pmyo-2::mCh]* | 106/120 | 69.98 | 2.52 | 1 | <0.0001 |
| **Trial 3^** |  |  |  |  |  |  |
| N2 | Wild type | 68 / 120 | 68.39 | 2.2 |  |  |
| CF1903 | *glp-1* | 102 / 117 | 94.91 | 3.2 | 0.0001 |  |
| AGP44 | *glp-1 glmEx10 [Pgly-19::nhr-49::GFP + Pmyo-2::mCh]* | 97 / 115 | 82.13 | 2.71 | 0.0044 | 0.0086 |

| Strain | Background/Genotype | Trial 1@ |  |  | Bonferroni P-value |  |
| --- | --- | --- | --- | --- | --- | --- |
|  |  | n = obs/<br>total | Mean (h) | SE +/- | P (vs N2) | P (vs *glp-1*) |
| AGP22 | *nhr-49;glp-1* | 72/72 | 11.27 | 0.26 |  | <0.0001 |
| CF1903 | *glp-1* | 65/77 | 25.78 | 0.98 |  |  |
| AGP44 | *glp-1 glmEx10 [Pgly-19::nhr-49::GFP + Pmyo-2::mCh]* | 26/27 | 20.06 | 1.09 |  | 0.0003 |
| **Trial 2~** |  |  |  |  |  |  |
| N2 | Wild type | 82 / 122 | 15.53 | 0.58 |  |  |
| CF1903 | *glp-1* | 112 / 124 | 22.98 | 0.67 | <0.0001 |  |
| AGP44 | *glp-1 glmEx10 [Pgly-19::nhr-49::GFP + Pmyo-2::mCh]* | 105 / 122 | 20.69 | 0.44 | <0.0001 | 0.0002 |
| **Trial 3\$** | | | | | | |
| N2 | Wild type | 73 / 121 | 17.81 | 0.56 |  |  |
| CF1903 | *glp-1* | 101 / 114 | 23.35 | 0.69 | <0.0001 |  |
| AGP44 | *glp-1 glmEx10 [Pgly-19::nhr-49::GFP + Pmyo-2::mCh]* | 101 / 116 | 20.93 | 0.41 | 0.0004 | 0.0004 |

\*, #, ^, &, @, ~ on different tabs/sub-sections of Table S6 denote trials performed at the same time Underlined Trials shown in Fig. 6

| Table S6D. 1: SURVIVAL ON PA14 |  |  |  |  |  |  |
| --- | --- | --- | --- | --- | --- | --- |
| Strain | Background/Genotype | Trial 1* |  |  | Bonferroni P-value |  |
|  |  | n = obs/<br>total | Mean (h) | SE +/- | P (vs N2) | P (vs<br>*glp-1*) |
| N2 | Wild type | 83 / 105 | 67.31 | 1.37 |  |  |
| CF1903 | *glp-1* | 116 / 127 | 83.68 | 2.04 | <0.0001 |  |
| AGP36 | *glp-1 glmEx7 [Pmyo-3::nhr-49::GFP + Pmyo-2::mCh]* | 96 / 118 | 73.05 | 2.12 | 0.0378 | 0.0107 |
| Trial 2# |  |  |  |  |  |  |
| N2 | Wild type | 61/112 | 72.21 | 3.63 |  |  |
| CF1903 | *glp-1* | 95/109 | 88.89 | 2.57 | 0.001 |  |
| AGP36 | *glp-1 glmEx7 [Pmyo-3::nhr-49::GFP + Pmyo-2::mCh]* | 110/120 | 80.21 | 2.84 | 0.3629 | 0.6544 |
| Trial 3^ |  |  |  |  |  |  |
| N2 | Wild type | 68 / 120 | 68.39 | 2.2 |  |  |
| CF1903 | *glp-1* | 102 / 117 | 94.91 | 3.2 | 0.0001 |  |
| AGP36 | *glp-1 glmEx7 [Pmyo-3::nhr-49::GFP + Pmyo-2::mCh]* | 110 / 120 | 88.62 | 2.93 | <0.0001 | 0.747 |
| Table S6D.2: SURVIVAL ON OP50 |  |  |  |  |  |  |
| Strain | Background/Genotype | Trial 1@ |  |  | Bonferroni P-value |  |
|  |  | n = obs/<br>total | Mean (h) | SE +/- | P (vs N2) | P (vs<br>*glp-1*) |
| AGP22 | *nhr-49;glp-1* | 72/72 | 11.27 | 0.26 |  | <0.0001 |
| CF1903 | *glp-1* | 65/77 | 25.78 | 0.98 |  |  |
| AGP36 | *glp-1 glmEx7 [Pmyo-3::nhr-49::GFP + Pmyo-2::mCh]* | 46/83 | 24.65 | 0.98 |  | 1 |
| Trial 2\$ | | | | | | |
| N2 | Wild type | 73 / 121 | 17.81 | 0.56 |  |  |
| CF1903 | *glp-1* | 101 / 114 | 23.35 | 0.69 | <0.0001 |  |
| AGP36 | *glp-1 glmEx7 [Pmyo-3::nhr-49::GFP + Pmyo-2::mCh]* | 104 / 121 | 21.89 | 0.55 | <0.0001 | 0.3937 |

\*, #, ^, &, @, ~ on different tabs/sub-sections of Table S6 denote trials performed at the same time Underlined Trials shown in Fig. 6

| Table S6E.1: SURVIVAL ON PA14 |  |  |  |  |  |  |
| --- | --- | --- | --- | --- | --- | --- |
| Strain | Background/Genotype | Trial 1* |  |  | Bonferroni P-value |  |
|  |  | n = obs/<br>total | Mean (h) | SE +/- | P (vs N2) | P (vs<br>*glp-1*) |
| N2 | Wild type | 83 / 105 | 67.31 | 1.37 |  |  |
| CF1903 | *glp-1* | 116 / 127 | 83.68 | 2.04 | <0.0001 |  |
| AGP41 | *glp-1 glmEx11 [Pcol-12::nhr-49::GFP + Pmyo-2::mCh]* | 80 / 98 | 69.92 | 2.58 | 0.0674 | 0.009 |
| Trial 2# |  |  |  |  |  |  |
| N2 | Wild type | 61/112 | 72.21 | 3.63 |  |  |
| CF1903 | *glp-1* | 95/109 | 88.89 | 2.57 | 0.001 |  |
| AGP36 | *glp-1 glmEx7 [Pmyo-3::nhr-49::GFP + Pmyo-2::mCh]* | 12./13 | 79.72 | 9.95 | 1 | 1 |
| Trial 3^ |  |  |  |  |  |  |
| N2 | Wild type | 68 / 120 | 68.39 | 2.2 |  |  |
| CF1903 | *glp-1* | 102 / 117 | 94.91 | 3.2 | 0.0001 |  |
| AGP41 | *glp-1 glmEx11 [Pcol-12::nhr-49::GFP + Pmyo-2::mCh]* | 47 / 60 | 65.17 | 4.78 | 1 | <0.0001 |
| Table S6E.2: SURVIVAL ON OP50 |  |  |  |  |  |  |
| Strain | Background/Genotype | Trial 1@ |  |  | Bonferroni P-value |  |
|  |  | n = obs/<br>total | Mean (h) | SE +/- | P (vs N2) | P (vs<br>*glp-1*) |
| AGP22 | *nhr-49;glp-1* | 72/72 | 11.27 | 0.26 |  | <0.0001 |
| CF1903 | *glp-1* | 65/77 | 25.78 | 0.98 |  |  |
| AGP41 | *glp-1 glmEx11 [Pcol-12::nhr-49::GFP + Pmyo-2::mCh]* | 28/42 | 24.06 | 0.83 |  | 0.0611 |
| Trial 2\$ | | | | | | |
| N2 | Wild type | 73 / 121 | 17.81 | 0.56 |  |  |
| CF1903 | *glp-1* | 101 / 114 | 23.35 | 0.69 | <0.0001 |  |
| AGP41 | *glp-1 glmEx11 [Pcol-12::nhr-49::GFP + Pmyo-2::mCh]* | 84 / 102 | 23.79 | 0.54 | <0.0001 | 1 |

\*, #, ^, &, @, ~ on different tabs/sub-sections of Table S6 denote trials performed at the same time Underlined Trials shown in Fig. 6

**Table S7:** Transgenic strains created for this study.

| Name | Genotype |
| --- | --- |
| N2 | Wild type, Ghazi Lab |
| CF1903 | *glp-1(e2144ts)* |
| AGP12a | *nhr-49(nr2041)*I Outcrossed Ghazi Lab N2 3x |
| AGP22 | *nhr-49(nr2041)* I; *glp-1(e2141ts)* III |
| AGP110 | *nhr-49(et7)* *gof* Outcrossed *nhr-49(et7);paqr-2* to Ghazi Lab N2 4x |
| VC1668 | *fmo-2(ok2147)* |
| RB1899 | *acs-2(ok2457)* |
| EB271 | *fmo-2p::gfp* |
| WBM170 | *wbmEx57 [acs-2p::GFP + rol-6(su1006)]* |
| **NHR-49 Transgenic Strains** |  |
| **Wild Type Background** |  |
| AGP24f | wild type; *glmEx5 [Pnhr-49::nhr-49::gfp + Pmyo-2::mCherry]* |
| AGP27a | wild type; *glmEx7 [Pmyo-3::nhr-49::GFP + Pmyo-2::mCherry]* |
| AGP40a | wild type; *glmEx9 [Pgly-19::nhr-49::GFP + Pmyo-2::mCherry]* |
| AGP45 | wild type; *glmEx11 [Pcol-12::nhr-49::GFP + Pmyo-2::mCherry]* |
| AGP104a | wild type; *glmEx21 [Prgef-1::nhr-49::GFP + Pmyo-2::mCherry]* |
| AGP102a | wild type; *glmEx20 [Punc-119::nhr-49::GFP + Pmyo-2::mCherry]* |
| ***nhr-49* Mutant Background** |  |
| AGP33a | *nhr-49(nr2041)*I;*glmEx5 [Pnhr-49::nhr-49::GFP + Pmyo-2::mCherry]* |
| AGP63 | *nhr-49(nr2041)*I; *glmEx8 [Pmyo-3::nhr-49::GFP + Pmyo-2::mCherry]* (25ng/uL) |
| AGP65 | *nhr-49(nr2041)*I; *glmEx9 [Pgly-19::nhr-49::GFP + Pmyo-2::mCherry]* |
| AGP53 | *nhr-49(nr2041)*I; *glmEx11 [Pcol-12::nhr-49::GFP + Pmyo-2::mCherry]* |
| AGP51 | *nhr-49(nr2041)*I; *glmEx13 [Prgef-1::nhr-49::GFP + Pmyo-2::mCherry]* |
| AGP103a | *nhr-49(nr2041)*I; *glmEx20 [Punc-119::nhr-49::GFP + Pmyo-2::mCherry]* |
| ***glp-1* Mutant Background** |  |
| AGP29d | *glp-1(e2141ts)* III;*glmEx6 [Pnhr-49::nhr-49::GFP + Pmyo-2::mCherry]* (25ng/uL).<br>Obtained by crossing AGP28a x CF1903. |
| AGP36a | *glp-1(e2141ts)* III; *glmEx7 [Pmyo-3::nhr-49::GFP + Pmyo-2::mCherry]* |
| AGP44 | *glp-1(e2141ts)* III; *glmEx10 [Pgly-19::nhr-49::GFP + Pmyo-2::mCherry]* (25ng/uL) |
| AGP41a/b | *glp-1(e2141ts)* III; *glmEx11 [Pcol-12::nhr-49::GFP + Pmyo-2::mCherry]* |
| AGP69 | *glp-1(e2141ts)* III; *glmEx13 [Prgef-1::nhr-49::GFP + Pmyo-2::mCherry]* |
| AGP105a/b | *glp-1(e2141)*; *glmEx20 [Punc-119::nhr-49::GFP + Pmyo-2::mCherry]* |
| ***nhr-49;glp-1* Mutant Background** |  |
| AGP34a | *nhr-49(nr2041)* I; *glp-1(e2141ts)* III; *glmEx5 [Pnhr-49::nhr-49::GFP + Pmyo-2::mCherry]* |
| AGP279 | *nhr-49(nr2041)* I; *glp-1(e2141ts)* III; *glmEx7 [Pmyo-3::nhr-49::GFP + Pmyo-2::mCherry]*. Obtained by crossing AGP36 x AGP12a |
| AGP43 | *nhr-49(nr2041);glp-1(e2141)*; *glmEx10 [Pgly-19::nhr-49::GFP + Pmyo-2::mCherry]* (25ng/uL) |
| AGP54a | *nhr-49(nr2041);glp-1(e2141)*; *glmEx11 [Pcol-12::nhr-49::GFP + Pmyo-2::mCherry]* |
| AGP70a/b | *nhr-49(nr2041);glp-1(e2141)*; AGP22;*glmEx13 [prgef-1::nhr-49::GFP + Pmyo-2::mCherry]* |
| AGP115a/b | *nhr-49(nr2041);glp-1(e2141)*; *glmEx29 [Punc-119::nhr-49::GFP + Pmyo-2::mCherry]* |

